# Exploiting protein language models for the precise classification of ion channels and ion transporters

**DOI:** 10.1101/2023.07.11.548644

**Authors:** Hamed Ghazikhani, Gregory Butler

## Abstract

This study presents TooT-PLM-ionCT, a composite framework consisting of three distinct systems, each with different architectures and trained on unique datasets. Each system within TooT-PLM-ionCT is dedicated to a specific task: segregating ion channels (ICs) and ion transporters (ITs) from other membrane proteins and differentiating ICs from ITs. These systems exploit the capabilities of six diverse Protein Language Models (PLMs) - ProtBERT, ProtBERT-BFD, ESM-1b, ESM-2 (650M parameters), and ESM-2 (15B parameters). As these proteins play a pivotal role in the regulation of ion movement across cellular membranes, they are integral to numerous biological processes and overall cellular vitality. To circumvent the costly and time-consuming nature of wet lab experiments, we harness the predictive prowess of PLMs, drawing parallels with techniques in natural language processing. Our strategy engages six classifiers, embracing both conventional methodologies and a deep learning model, for each of our defined tasks. Furthermore, we delve into critical factors influencing our tasks, including the implications of dataset balancing, the effect of frozen versus fine-tuned PLM representations, and the potential variance between half and full precision floating-point computations. Our empirical results showcase superior performance in distinguishing ITs from other membrane proteins and differentiating ICs from ITs, while the task of discriminating ICs from other membrane proteins exhibits results commensurate with the current state-of-the-art.

**Author summary:** In our research, we have designed TooT-PLM-ionCT, a composite framework composed of three unique systems, each tailored to a specific protein classification task and trained on different datasets. This framework is our tool for categorizing integral membrane proteins, specifically ion channels and ion transporters. These proteins are essential to the health of cells, as they manage ion movement across cell membranes. To bypass the high costs and long timelines of conventional lab experiments, we have turned to advanced computation methods akin to how computers process human language. Our three-pronged approach harnesses six top-tier Protein Language Models and a range of classifiers to discern between these key proteins. In doing so, we also evaluated the effects of various conditions, like dataset balance, representation methods, and levels of computation precision, on the accuracy of our classification tasks. The outcomes show our framework effectively identifies ion transporters, sets them apart from ion channels, and distinguishes ion channels on par with existing top-notch techniques. The performance, however, can vary based on the task, suggesting that customizing the approach for each task could be beneficial. In the future, we plan to expand the depth and breadth of our protein study by incorporating additional knowledge sources, utilizing more refined representation methods, and testing our framework on larger and diverse protein datasets. This progress sets us on a path to better understand proteins and their roles in cellular health.

## Introduction

### Background

Protein language models (PLMs) are a transformative development in the field of bioinformatics, leveraging the power of machine learning to predict protein structures and functions from their amino acid sequences [1–3]. These models, inspired by natural language processing (NLP) techniques [4–7], treat proteins as “sentences” composed of “words” (amino acids), enabling the prediction of protein properties based on sequence information alone [8]. The importance of PLMs lies in their potential to revolutionize our understanding of proteins, the building blocks of life, and to accelerate drug discovery and design processes [9]. They provide a powerful tool for predicting protein structures, which is crucial for understanding diseases and developing treatments [10]. Moreover, PLMs produce comprehensive representations of protein sequences that are useful for various applications in protein analysis, including predicting protein function, protein-protein interactions, and protein structure [1, 3,11–17]. Unsal et al. [8] review the use of natural language models for protein representation from 2015 to the present.

The regulation of ion movement across cell membranes is a critical aspect of cellular function, with ion channels (ICs) and ion transporters (ITs) playing key roles [18]. These membrane proteins (MPs) are involved in maintaining ion homeostasis (the regulation and maintenance of a stable and balanced concentration of ions), regulating transmembrane potential, and facilitating electrical signaling, which are essential for various cellular processes such as proliferation, migration, apoptosis, and differentiation [19–21].

ITs, also known as ion pumps, actively transport ions against their concentration gradient, a process that requires potential energy [22]. On the other hand, ICs are transmembrane protein complexes located in the lipid bilayer membrane of all cells [23]. They facilitate the passive movement of ions across cell membranes, thereby helping cells maintain electrical properties and regulate functions [19, 20].

Given their crucial role in cellular function, ICs have become a significant focus in membrane protein research and drug discovery [23]. They serve as promising therapeutic targets for various diseases, including neurological disorders, cardiovascular diseases, and cancer [24–27].

In an effort to expedite the drug discovery process and circumvent the high costs and time-consuming nature of wet lab experiments, computational methods have been developed. These innovative techniques efficiently predict the presence and function of ion channels, thereby accelerating the identification of potential drug targets [28, 29].

Among these computational methods, PLMs have emerged as a particularly powerful tool [3]. By learning the sequence patterns of different protein families, PLMs can accurately classify proteins and predict their functions [17, 23, 30]. This capability not only streamlines the process of protein classification but also opens up new avenues for the discovery of therapeutic targets [31].

### Review of previous work

There has been a significant amount of research on predicting ICs and ITs in the past, with an emphasis on developing computational methods that can accurately differentiate these proteins from other MPs [22, 28, 29, 32–35]. These methods have often utilized traditional machine learning techniques, such as Support Vector Machines (SVM) and Random Forests (RF), which classify protein sequences based on features derived from their primary, secondary, and tertiary structures. These features can include information about the sequence itself, such as the presence of certain amino acid residues or motifs, as well as structural features, such as secondary structure elements or solvent accessibility [29, 36]. The use of these features for ion channel prediction is thoroughly explained in Menke et al. [29] and Ashrafuzzaman [28].

The advent of deep learning has paved the way for novel opportunities in predicting ICs and ITs. Recent studies underscore the potential of these advanced techniques to generate intricate representations of protein sequences, thereby enhancing the efficiency of IC and IT prediction models [22, 35]. In their respective methodologies, Taju and Ou [35], as well as Nguyen et al. [22], utilized position-specific scoring matrices (PSSM) for encoding proteins into feature vectors, while leveraging Convolutional Neural Networks (CNNs) for classifying ICs and ITs from other membrane proteins (MPs). These innovative models could discern complex patterns in protein sequences, employing this information to augment prediction performance, potentially surpassing the constraints of conventional machine learning approaches [22, 35]. However, it is noteworthy that their work primarily focuses on distinguishing ion channels from other membrane proteins and ion transporters from other membrane proteins, rather than the task of differentiating ion channels from ion transporters.

Ghazikhani et al. pioneered the introduction of TooT-BERT-T [30] and TooT-BERT-C [23], sophisticated methods designed for distinguishing transmembrane transport proteins from non-transport proteins, as well as differentiating ICs from non-ICs. These methods incorporate a Logistic Regression (LR) classifier with fine-tuned representations derived from a PLM known as ProtBERT-BFD [3]. As the most advanced predictors for transporters and ICs, these approaches underscore the promising potential of employing protein language models for such tasks.

### Research overview and objective

In this comprehensive study, we introduce TooT-PLM-ionCT, a multi-faceted framework comprised of three distinct systems, each designed to effectively distinguish ion channels, ion transporters, and other membrane proteins. These systems are tailored to their respective tasks and trained on different datasets, enabling precise classification. As part of our research, we also rigorously analyze six PLMs in conjunction with six different classifiers. Additionally, we delve into key variables influencing PLM performance in protein classification tasks, such as dataset balancing, representation tuning, and the precision of floating-point calculations.

The primary aim of this paper is to present a trailblazing, automated approach that precisely categorizes ion transporters and ion channels within the vast spectrum of membrane proteins. By shedding light on the intricate nature of these crucial biological elements, our goal is to simplify their identification in bioinformatics research. This could potentially expedite the discovery of new therapeutic targets for a plethora of diseases, highlighting the substantial potential of our approach.

### Study of impacts

In this study, we embarked on a meticulous investigation of three pivotal factors that could significantly influence the performance of PLMs in our tasks. These encompass:

- The choice between using frozen or fine-tuned PLM representations.
- The influence of balanced versus imbalanced datasets on model performance.
- The implications of half-precision versus full-precision floating-point computations.

Each of these elements represents a vital facet of the model’s configuration and data management, thus underscoring the importance of their potential impacts on model performance. The forthcoming sections deliver a succinct synopsis of each factor, explicating the fundamental concept and the rationale for its incorporation in our study.

#### Frozen vs fine-tuned representations

The concept of frozen and fine-tuned representations pertains to the degree of adaptation of pre-trained language models to a specific task. Frozen representations refer to the utilization of pre-trained models in their original state, without any further task-specific training. On the other hand, fine-tuned representations involve the additional step of task-specific training, where the pre-existing parameters of the pre-trained models are adjusted to enhance their performance on the given task.

The comparative study of frozen and fine-tuned versions of a PLM offers valuable insights into the performance dynamics of these models. It allows us to understand the inherent behavior of the original pre-trained models (as reflected in the frozen state) and to quantify the extent of improvement achievable through task-specific fine-tuning. This comparison can potentially expose the limitations of the pre-training process and highlight the areas where fine-tuning can yield significant benefits.

It is important to note that fine-tuning necessitates additional computational resources compared to the use of frozen models. Consequently, if the performance enhancement achieved through fine-tuning is marginal or negligible for a specific task, it might be more resource-efficient to employ the model in its frozen state. This aspect underscores the importance of our investigation into the relative merits of frozen and fine-tuned representations in the context of our tasks.

#### Balanced vs imbalanced datasets

The terms “balanced” and “imbalanced” in machine learning refer to the distribution of classes within a dataset. A balanced dataset exhibits approximately equal representation of all classes, while an imbalanced dataset is characterized by unequal representation of classes. In the context of this study, these terms are used to describe the distribution of membrane protein sequences in the DS-C dataset (Table 2).

Imbalanced datasets, where certain classes are underrepresented, can significantly impact the performance of a machine learning model. The model may develop a bias towards the majority class, leading to suboptimal performance when predicting the minority class. In the realm of PLMs, this issue translates into a potential struggle for the model to accurately predict protein types that are underrepresented in the training data.

Furthermore, the bias introduced by an imbalanced dataset can result in a model that performs better for the class with greater representation in the data. For instance, if the dataset contains a significantly larger number of MPs compared to ICs or ITs, the model may develop a bias towards MPs. This bias could compromise the model’s ability to accurately predict ICs or ITs, underscoring the importance of considering the balance of classes in the dataset.

#### Half vs full precision floating points calculations

Half and full precision floating-point representations pertain to the level of numerical precision employed in model computations. Full precision, typically realized through 32-bit floats, provides superior numerical accuracy. Conversely, half precision, utilizing 16-bit floats, curtails memory usage and computational demands, albeit at the expense of a slight reduction in numerical accuracy.

The use of half-precision computations can expedite the training process, but it may also influence model performance due to the diminished numerical precision. It is crucial to evaluate whether this reduction in precision significantly affects the model’s capacity to learn and generalize effectively.

Additionally, investigating the impact of half versus full precision provides valuable insights into the balance between computational efficiency and model performance. This understanding facilitates informed decision-making, taking into account the available computational resources and the precision requirements of the task at hand.

### Paper structure

This paper is organized as follows: Section details our methodologies, including the datasets used and the process for balancing the membrane proteins dataset. It provides a brief overview of the employed PLMs and classifiers, elaborates on hyperparameter optimization, and discusses the evaluation metrics used to assess model performance. In Section, we present and dissect the results of our experimental analyses. This section evaluates the performance of different PLMs and classifiers for each task, sheds light on the impact of the three previously mentioned factors, and includes visualizations of protein representations. Additionally, it juxtaposes our findings with current state-of-the-art methodologies for each task. Finally, Section encapsulates our contributions and the insights gleaned from our study. It also outlines potential future research avenues, emphasizing areas where additional exploration could enrich the understanding of protein classification using PLMs.

## Materials and methods

### Methodology overview

We have undertaken a comprehensive evaluation of representations derived from six distinct PLMs. These include ProtBERT, ProtBERT-BFD, and ProtT5 from ProtTrans project [3], as well as ESM-1b, ESM-2, and ESM-2 15B from ESM project [2, 37].

To further our analysis, we have employed six classifiers with the aim of distinguishing ICs from other MPs, differentiating ITs from other MPs, and discriminating ICs from ITs. These classifiers encompass traditional methodologies such as LR, k-Nearest Neighbor (kNN), support vector machine (SVM), random forest (RF), and feed-forward neural network (FFNN). Additionally, we have incorporated a convolutional neural network (CNN), a deep learning model, for comparative analysis.

Our study also delves into the examination of several critical factors that could potentially influence the outcomes of our tasks. These include the impact of balancing the MP dataset on the results, the influence of frozen and fine-tuned representations from PLMs, and the potential differences between half and full precision floating-point calculations. By investigating these factors, we aim to provide a more nuanced understanding of the performance and applicability of PLMs in protein classification tasks. Refer to Table 1 for a comprehensive summary of the research methodology employed in this study.

**Table 1.** Comprehensive overview of research methodology. This table encapsulates the various components of the research methodology employed in this study, providing a concise summary and brief description of each element.

| Methodology Component | Details |
| --- | --- |
| Protein Language Models | ProtBERT, ProtBERT-BFD, ProtT5 (ProtTrans project), ESM-1b, ESM-2, ESM-2.15B (ESM project) |
| Tasks | Discrimination of ion channels vs other membrane proteins, ion transporters vs other membrane proteins, ion channels vs ion transporters |
| Classifiers | SVM, Logistic Regression (LR), Random Forest (RF), kNN, Feed-forward Neural Network (FFNN), CNN |
| Hyperparameter Optimization | Grid search using scikit-learn (for SVM, LR, RF, kNN, FFNN) and Optuna (for CNN) |
| Cross-Validation Technique | 5-fold cross-validation |
| Evaluation Metrics | Accuracy, MCC, Sensitivity, Specificity |
| Statistical Significance Analysis | Paired Student t-test, ANOVA |
| Impacts Evaluated | 1) Frozen vs fine-tuned representations from PLMs, 2) Balanced vs imbalanced datasets (Downsampling of MPs dataset), 3) Half vs full precision floating point calculations |
| Presentation of Results | Tables and figures, grouped results by various aspects such as dataset balance, classifier type, PLM, representation type (frozen or fine-tuned), precision type (half or full), and UMAP projection figures for each PLM, task, and representation type |
| Optimal Configuration | Selected the best configuration for each task, ran on independent test set, and compared results with state-of-the-art |
| Limitations | Could not fine-tune large PLMs like ProtT5 and ESM-2.15B due to resource constraints (GPUs, memory), could not extract full precision floating point from these PLMs. This led to missing values in tables and figures. |

### Dataset

In our study, we employ the same dataset used in the DeepIon [35] and MFPS CNN [22] projects, which was gathered from the UniProt database [38]. To ensure a diverse and representative collection, Taju and Ou [35] applied the BLAST algorithm [39] to remove protein sequences with more than 20% similarity. The resulting dataset comprises 4915 protein sequences, including 301 ion channels, 351 ion transporters, and 4263 (other) membrane proteins. The dataset was split into training and test sets for assessing model generalizability. The distribution of sequences in the dataset is presented in Table 2.

**Table 2.**
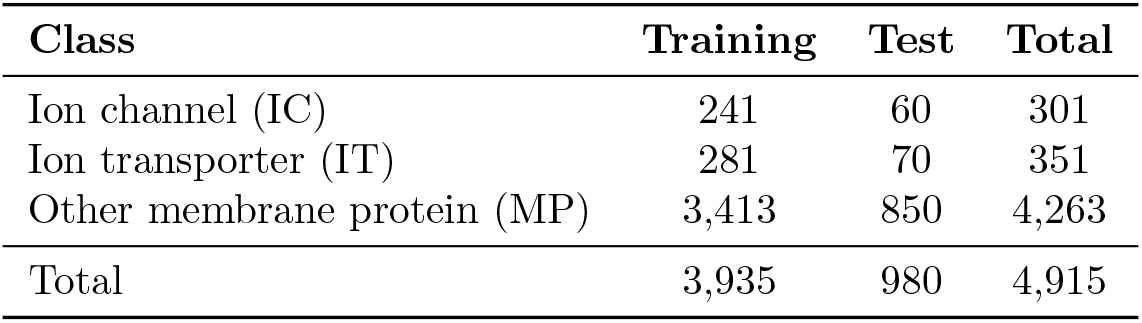
DS-C, the ion channel and ion transporter dataset. This table displays the distribution of sequences in the dataset used in this study, separated into the training and test sets.

| Class | Training | Test | Total |
| --- | --- | --- | --- |
| Ion channel (IC) | 241 | 60 | 301 |
| Ion transporter (IT) | 281 | 70 | 351 |
| Other membrane protein (MP) | 3,413 | 850 | 4,263 |
| Total | 3,935 | 980 | 4,915 |

#### Balancing the membrane protein dataset

As highlighted in Table 2, there exists a significant disparity in the number of membrane protein sequences in comparison to ion channel or ion transporter protein sequences. For this study, our objective was to assess the performance of PLMs and classifiers employing both imbalanced and balanced datasets. To construct a balanced dataset (Fig 1), we implemented a random selection process to draw 280 sequences from the membrane protein training set. To enhance the accuracy of the results and mitigate potential variability, this process was reiterated ten times, each iteration using a distinct random state.

**Fig 1.**
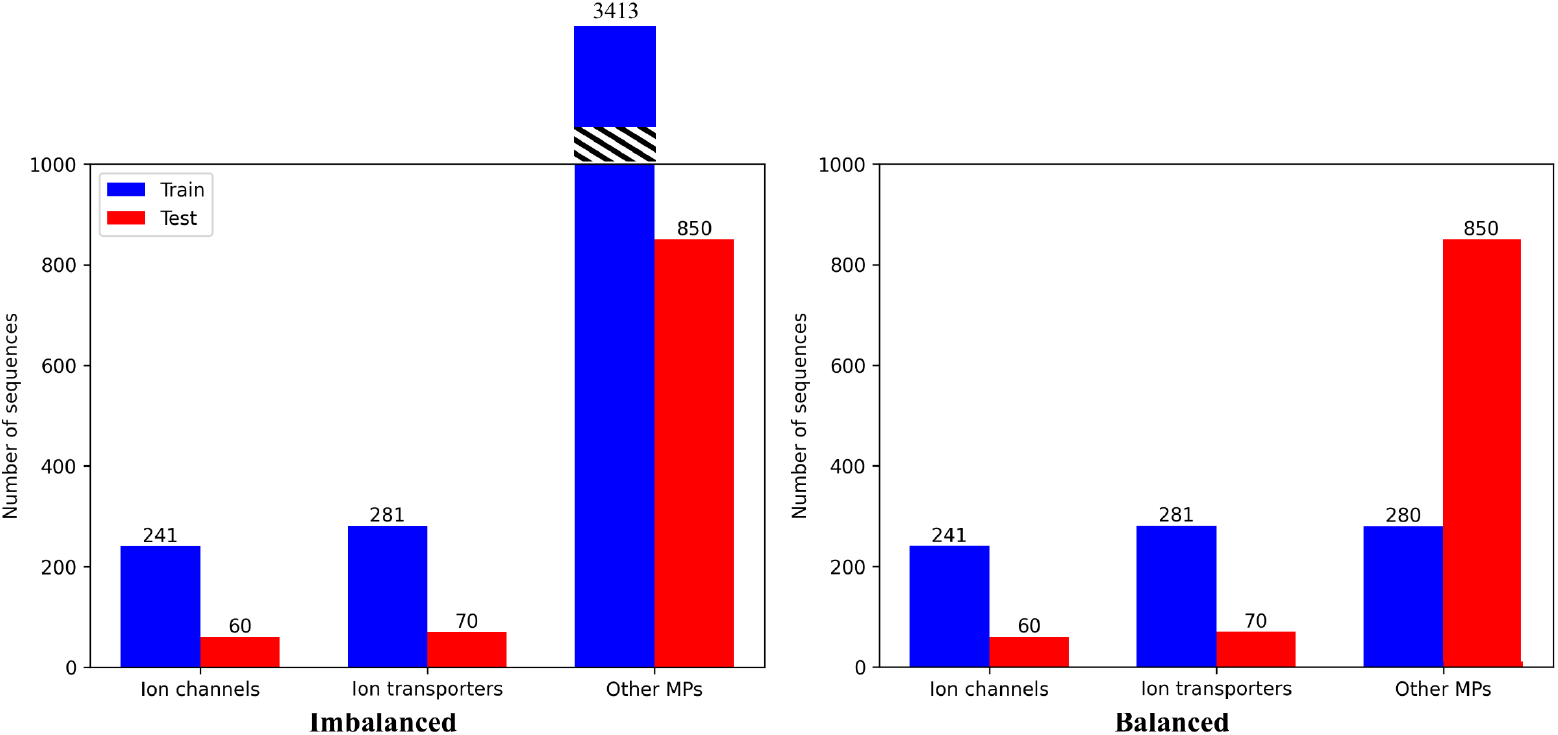
Visualization of Membrane Protein Dataset Balancing. This figure presents the distribution of sequences in each dataset, delineated as bar plots. The training set sequences are represented by the blue bars, whereas the red bars depict the sequences in the independent test set. The left-hand figure portrays the distribution within the imbalanced dataset of additional membrane proteins (MPs). Conversely, the right-hand figure exhibits the balanced dataset, which was achieved through undersampling of MPs in the training set.

### Protein language models (PLMs)

This study leverages six distinct Protein Language Models (PLMs) for comparative analysis (Table 3): (1) *ProtBERT* [3] is an encoder-only model inspired by BERT [40], pre-trained on UniRef100 [41]. (2) *ProtBERT-BFD* [3], analogous to ProtBERT, is pre-trained on the BFD database [42] instead of UniRef100. (3) *ProtT5-XL* [3] (simplified to *ProtT5* for convenience), is an encoder-decoder model rooted in the T5 architecture [6]. It is initially trained on BFD and subsequently fine-tuned on Uniref50 [41]. (4) *ESM-1b* [2] is a Transformer model pre-trained on UniRef50. (5) *ESM-2* [37], while akin to ESM-1b, benefits from enhanced architecture, improved training parameters, and augmented computational resources and data. (6) *ESM-2 15B* [37], the largest PLM to date, is a more extensive version of ESM-2, incorporating 15 billion parameters.

**Table 3.** Comparative analysis of various PLMs utilized in this study. This table provides a detailed comparison of PLMs used in this research, including ProtBERT [3], ProtBERT-BFD [3], ProtT5 [3], ESM-1b [2], ESM-2 [37], and ESM-2 15B [37].

|  | ProtBERT | ProtBERT-BFD | ProtT5 | ESM-1b | ESM-2 | ESM-2.15 |
| --- | --- | --- | --- | --- | --- | --- |
| Parameters | 420M | 420M | 3B | 650M | 650M | 15B |
| Dataset | UniRef100 | BFD | BFD | UniRef50 | UniRef50 | UniRef50 |
| Sequences | 216M | 2.1B | 2.1B | 27M | 27M | 27M |
| Embedding dim | 1024 | 1024 | 1024 | 1280 | 1280 | 5120 |
| Layers | 30 | 30 | 24 | 33 | 33 | 48 |

To derive frozen representations, we harness feature vectors from the final layer of the PLMs, employing mean-pooling to generate a unique representation for each protein sequence. This process is consistent with the methodologies adopted in ProtTrans [3] and ESM [2, 37].

For fine-tuning of the PLMs, we engage the Trainer API from the transformers library [43]. We primarily utilize the library’s default hyperparameters but modify the number of epochs to 5, following the guidelines of the original BERT paper [40]. To mitigate memory constraints, we adopt a batch size of 1.

### Classifiers

For our machine learning classifiers, we implement Support Vector Machine (SVM) [44], k-Nearest Neighbors (kNN) [45], Random Forest (RF) [46], Feed-Forward Neural Network (FFNN) [47], and Logistic Regression (LR) [48] using the scikit-learn library [49], whereas Convolutional Neural Network (CNN) [50] using PyTorch [51]. These classifiers are designed to provide a comprehensive comparison of various machine learning approaches in combination with the PLMs.

### Hyperparameter optimization

In this investigation, we incorporated an all-encompassing strategy for hyperparameter optimization, harnessing the prowess of scikit-learn grid search [49] and Optuna [52], an advanced Python library specifically designed for hyperparameter optimization. The primary objective was to discern the quintessential set of hyperparameters for each model to maximize the efficacy of our classification algorithms.

With respect to conventional classifiers such as SVM, RF, kNN, LR, and FFNN, we exploited grid search—an exhaustive technique that systematically scrutinizes a pre-defined subset of hyperparameters. This process was executed utilizing the scikit-learn library [49].

Each classifier was assigned a unique set of hyperparameters to investigate. The specific grids of hyperparameters tailored for each classifier were as follows:

- SVM: The investigation included cost parameters (C) of 0.1, 1, 10, and 100; kernel coefficients (gamma) of 0.1, 1, and 10; and kernel types (kernel) inclusive of linear, rbf, and sigmoid.
- RF: The search encompassed the number of trees in the forest (number of estimators) of 50, 100, and 200; the maximum tree depth (maximum depth) of 5, 10, and None; and the minimum samples required to split an internal node (minimum samples split) of 2, 5, and 10.
- kNN: The evaluation incorporated the number of considered neighbors (number of neighbors) of 3, 5, 7, and 9; the prediction weight function (weights) of uniform and distance; and the algorithm used for calculating the nearest neighbors (algorithm) of ball tree, kd tree, and brute.
- LR: The investigation comprised various penalty types (penalty) of l1 and l2; cost parameters (C) of 0.1, 1, 10, and 100; and optimization solvers (solver) of liblinear and saga.
- Feed-Forward Neural Network (FFNN): The search included the number of neurons in the hidden layer (hidden layer sizes) of (512, 256, 64), (512,), and (256,); the activation function for the hidden layer (activation) of relu and tanh; and the weight optimization solver (solver) of adam and sgd.

For the evaluation of model performance for each hyperparameter combination, we employed stratified 5-fold cross-validation. The optimization scoring metric was the Matthews Correlation Coefficient (MCC).

In the case of our Convolutional Neural Network (CNN) model, we utilized Optuna [52], a Python library adept at hyperparameter optimization. Optuna leverages a variety of optimization algorithms to traverse the hyperparameter space with the goal of identifying the optimal values that enhance the model’s performance.

The optimization procedure was encapsulated in an objective function, which incorporated the hyperparameters to be optimized. The specific hyperparameters and their respective ranges or sets of values were as follows:

- Kernel Sizes: The possibilities included combinations of [3, 5, 7], [3, 7, 9], [5, 7], and [7, 7, 7].
- Output Channels: The combinations were [128, 64, 32].
- Dropout Probability: The range was set from 0.2 to 0.5.
- Optimizer: The options included Adam, RMSprop, and SGD.
- Learning Rate: The range extended from 1e-6 to 1e-2 on a logarithmic scale.

The model underwent training for 10 epochs, with the performance being assessed on the validation set using MCC as the performance metric. The pruning feature of Optuna was harnessed to curtail trials early if they lacked promise, thereby conserving computational resources.

Owing to the intensive computational requirements of this procedure in terms of time and memory, the optimization was carried out singularly for each task and dataset, thereby resulting in five distinct hyperparameter settings (IC-MP balanced, IC-MP imbalanced, IT-MP balanced, IT-MP imbalanced, and IC-IT). For balanced datasets, one dataset was randomly selected from a pool of 10 for consideration.

The optimization procedure was executed for 100 trials, with each trial embodying a complete execution of the objective function with a distinct set of hyperparameters. The Optuna study was configured to maximize the MCC, and the optimization procedure was expedited by using a GPU for increased efficiency.

### Cross-validation and evaluation metrics

The technique of k-fold cross-validation, ubiquitously utilized in model evaluation, necessitates partitioning the original dataset into k subsets or folds of equivalent size. During each iteration, a single fold is reserved for validation, while the remaining k-1 folds serve as the training set. This cycle is repeated k times, ensuring each fold is used precisely once as the validation set. The model’s performance is then evaluated as the mean over the k iterations, delivering a more robust and accurate assessment of its capability to generalize. k-fold cross-validation plays a pivotal role in mitigating overfitting risk and curtailing bias in model evaluation. Our experimentation was conducted using 5-fold cross-validation, signifying the partitioning of the dataset into five subsets and repeated model training and validation over five iterations, with each fold serving as the validation set once.

In the context of this paper, we utilized four performance metrics to assess the efficacy of our approach for the tripartite tasks of IC-MP, IT-MP, and IC-IT. These metrics encompassed MCC, Accuracy, Sensitivity, and Specificity.

Accuracy represents an overall measure of correct classification rate, computed as the fraction of correct predictions relative to the total number of predictions. It is expressed as a percentage and can be determined using the following formula:

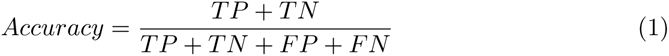

Sensitivity, also referred to as the true positive rate, measures the proportion of actual positive instances that are correctly identified. Its calculation is as follows:

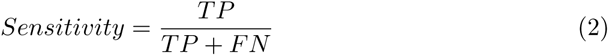

Specificity, alternatively known as the true negative rate, quantifies the proportion of actual negative instances that are correctly identified. Its calculation is as follows:

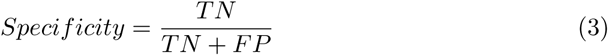

MCC is esteemed as a reliable and stable evaluation metric when handling imbalanced data [53]. The MCC values span from -1 to 1, where 1 signifies perfect prediction, 0 denotes performance equivalent to random chance, and -1 represents a total misalignment between predictions and observations. A high MCC value suggests a predictor demonstrating high accuracy for both positive and negative classes while maintaining a low misprediction rate for each class. In our research, we accord greater emphasis to the MCC metric due to its comprehensive nature and reliability.

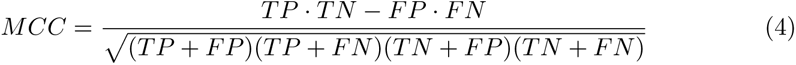

Here, TP (True Positive) denotes an instance where the classifier accurately predicts the positive class, TN (True Negative) signifies an instance where the classifier accurately predicts the negative class, FP (False Positive) represents an instance where the classifier erroneously predicts the positive class, and FN (False Negative) refers to an instance where the classifier inaccurately predicts the negative class.

### Statistical significance analysis

The statistical significance of observed differences was tested using two methods: the paired Student’s t-test [54] and Analysis of Variance (ANOVA) [55]. The paired Student’s t-test, ideal for comparing means of two related groups, was employed for two sets of related observations. Conversely, ANOVA, which assesses means across three or more unrelated groups, was applied when more than two independent groups were to be compared. The outcomes were expressed as a p-value, a statistical measure estimating the probability of random chance producing the observed results. Conventionally, a p-value below 0.05 signifies statistical significance, indicating a minimal probability that the observed difference occurred due to random chance. In our analysis, the p-value was computed from MCC metric, which is deemed comprehensive and reliable.

### Limitation

Our study was not without its limitations, primarily due to the constraints imposed by the available computational resources. The fine-tuning of large-scale PLMs such as ProtT5 (with 3 billion parameters) and ESM-2 15B (with 15 billion parameters) necessitates substantial computational resources and significant GPU memory, particularly for the extraction of full-precision floating-point representations [56]. Given our limited resources, which included a single GPU V100, we were unable to perform these tasks, resulting in some missing results in Section in our tables and figures.

Furthermore, the absence of results in Table 10 (ion channels vs ion transporters) is attributed to the fact that the corresponding studies [22, 35] do not report these specific results, and there are no readily available tools that can generate them. The primary focus of these papers is to classify ion channels and ion transporters against other membrane proteins, rather than against each other. However, in light of the data available to us, we chose to conduct this experiment and compare our models in this context as well.

## Results and discussion

This section presents a comprehensive exploration of the findings derived from our research, articulated through a combination of tables and figures to demonstrate and contrast varying facets of the study. We elucidate the performance of six distinct Protein Language Models (PLMs) as they engage with three specific tasks: differentiating ion channels (IC) from membrane proteins (MP), distinguishing ion transporters (IT) from MPs, and discerning IC from IT. We delve into the performance of six classifiers within these tasks, shedding light on three pivotal factors under investigation: the influence of frozen versus fine-tuned representations, the effect of balanced versus imbalanced datasets, and the impact of half versus full precision floating-point calculations.

Our findings are quantified using four evaluative metrics: Matthews Correlation Coefficient (MCC), Accuracy, Sensitivity, and Specificity. We present these results as mean ± standard deviation, obtained from a 5-fold cross-validation (CV). In our attempt to provide an overarching view, we compute averages over tasks, PLMs and classifiers, yielding a high-level depiction of our results. It should be noted, however, that results compared against the state-of-the-art are derived from an independent test set, employed solely for this purpose, with all other evaluations conducted on the training set.

In our tables, the highest values for each column and category are highlighted in bold, facilitating immediate recognition. Where there are more than two comparable values, the second highest are underlined to illustrate the proximity between the best and second-best results. In the corresponding figures, we prioritize the MCC metric, owing to its reliability and comprehensive nature. Each bar in these figures denotes the mean MCC, with the error bar atop indicating the standard deviation from the 5-fold CV. A Δ symbol highlights the difference between pairs of bars.

To ascertain the statistical significance of our findings, we employ ANOVA [55], a method for comparing the means of three or more groups, and the paired t-test [54], used to compare the means of two related groups. A p-value of 0.05 or smaller indicates a significant difference. It is important to note that this section primarily discusses general findings; more detailed results can be found in the appendix of this paper.

### Performance of PLMs for classification tasks

Table 4 presents a detailed evaluation of the six PLMs engaged in three distinct classification tasks: differentiating IC from MP, distinguishing IT from MP, and discerning IC from IT.

**Table 4.** Performance overview of protein language models for protein classification tasks. This figure provides a comprehensive performance evaluation of various protein language models (PLMs), organized in the order of their parameter count, across three distinctive protein classification tasks: differentiating ion channels (IC) from membrane proteins (MP), distinguishing ion transporters (IT) from MPs, and discerning IC from IT. The evaluation metrics, captured through a 5-fold cross-validation approach, are presented as mean±standard deviation. The p-value accompanying each result measures the statistical significance of observed differences among the PLMs. The highest value achieved for each task and column is highlighted in bold, whereas the second highest value is underlined to allow for comparative analysis between top-performing models.

| Task | PLM | MCC | Accuracy | Sensitivity | Specificity | P-value |
| --- | --- | --- | --- | --- | --- | --- |
| IC-MP | ProtBERT | 0.73±0.05 | 90.99±1.76 | 76.88±4.89 | 91.69±2.83 | 1.25e-06 |
|  | ProtBERT-BFD | 0.74±0.05 | 91.46±1.63 | 76.18±4.82 | 92.27±2.60 |  |
|  | ESM-1b | **0.84±0.03** | **94.15±1.17** | **88.44±3.39** | **94.33±1.91** |  |
|  | ESM-2 | <u>0.83±0.04</u> | <u>93.89±1.27</u> | <u>85.66±4.43</u> | **94.39±1.94** |  |
|  | ProtT5 | 0.79±0.05 | <u>93.12±1.38</u> | 79.68±4.98 | **94.35±1.81** |  |
|  | ESM-2.15B | 0.78±0.04 | <u>93.16±1.23</u> | 81.52±4.38 | <u>93.13±1.71</u> |  |
| IT-MP | ProtBERT | 0.71±0.05 | 90.75±1.41 | 75.66±4.69 | 91.58±2.34 | 2.49e-03 |
|  | ProtBERT-BFD | 0.74±0.05 | 91.10±1.64 | 78.91±4.79 | 92.30±2.33 |  |
|  | ESM-1b | **0.82±0.04** | **93.47±1.31** | **85.09±3.46** | **94.53±2.09** |  |
|  | ESM-2 | <u>0.78±0.04</u> | <u>92.64±1.36</u> | <u>82.06±4.20</u> | <u>93.41±2.26</u> |  |
|  | ProtT5 | 0.75±0.04 | <u>92.78±1.13</u> | 77.55±4.42 | <u>93.58±1.94</u> |  |
|  | ESM-2.15B | 0.72±0.04 | 91.58±1.46 | 76.12±4.26 | 91.90±2.32 |  |
| IC-IT | ProtBERT | 0.79±0.03 | 89.33±1.67 | 88.92±4.38 | 89.62±4.46 | 2.14e-06 |
|  | ProtBERT-BFD | 0.78±0.05 | 88.71±2.46 | 88.29±5.12 | 89.29±4.67 |  |
|  | ESM-1b | **0.85±0.04** | **92.46±2.25** | **92.83±3.42** | **92.12±4.21** |  |
|  | ESM-2 | 0.83±0.04 | 91.42±2.17 | 91.21±3.62 | 91.83±4.21 |  |
|  | ProtT5 | 0.84±0.04 | 91.83±1.83 | 91.00±2.67 | **92.50±3.83** |  |
|  | ESM-2.15B | **0.85±0.03** | **92.33±1.67** | <u>91.50±2.67</u> | **92.83±3.83** |  |

#### Performance of PLMs

Our findings underscore the superior performance of the ESM-1b PLM, as it eclipses other PLMs across all evaluation metrics and tasks. The lone exception is observed in the task of distinguishing IC from IT, where ESM-1b shares the lead position with ESM-2 15B. This indicates that ESM-1b consistently delivers high accuracy in predicting ICs and ITs from MPs.

However, the second-best performing model varies according to the task. ESM-2 exhibits commendable performance for differentiating IC from MP and distinguishing IT from MP, whereas ProtT5 excels in the IC-IT classification task. The significant variations in p-values across all PLMs further accentuate the formidable performance of ESM-1b.

In tasks pertaining to the differentiation of IC from MP and the distinction between IC and IT, the performance variance between the highest-ranking and the runner-up PLMs is minimally noticeable across all evaluation metrics. However, when tasked with discerning IT from MP, a notable performance discrepancy becomes apparent, particularly evident in the Matthews correlation coefficient (MCC) metric. This highlights a more substantial divergence in the proficiency of the two leading models, specifically ESM-2 and its predecessor, ESM-1b, within this particular task.

##### Factors contributing to ESM-1b’s superior performance outcomes

Our study posits that the unique architectural design of ESM-1b substantially contributes to its superior performance. This hypothesis is supported by our observation that identical pretraining dataset sizes, as employed in ESM-1b, ESM-2, ESM-2 15B, and more data in ProtBERT, ProtBERT-BFD, and ProtT5, in conjunction with model dimensions varying from 650M (for ESM-1b) to 15B (for ESM-2 15B) parameters, does not affect the performance of the corresponding PLMs significantly. We attribute the performance differences primarily to two factors: positional encoding and dropout strategies.

##### Positional encoding and its impact

ESM-1b [2] exhibits a unique approach to positional encoding. Diverging from the original Transformer architecture [4], it replaces the conventional static sinusoidal encoding with a learned encoding approach. This is markedly different from the approaches observed in the ESM-2 [37] and ProtTrans PLM [3] families.

##### Dropout strategies and their influence

Dropout [57], a prominent regularization technique in deep learning, randomly disables certain neural network units during training. This strategy enforces the network to develop more robust and generalizable features by reducing overfitting.

Distinct dropout strategies underscore a significant differentiation between ESM-1b and other PLMs. For instance, ESM-2 chooses to completely forgo dropout within hidden layers and attention. This pattern is also discernible in ProtBERT and ProtBERT-BFD, where dropout appears to be absent from their architectures. Conversely, ESM-1b not only incorporates dropout in its architectural framework but also applies it across various tasks. Considering the potential benefits of overfitting prevention measures, especially pertinent to the tasks investigated in our study, this difference assumes substantial significance.

Thus, in light of these findings, we suggest that the distinctive architectural design of ESM-1b plays a crucial role in facilitating its superior performance outcomes.

#### Impact of dataset balance and fine-tuning

This study observes that larger models, namely ProtT5 and ESM-2 15B, despite being precluded from fine-tuning due to resource constraints, managed to equal the performance of the smaller model, ESM-1b, on the balanced IC-IT dataset. Intriguingly, even with the application of fine-tuning to ESM-1b, the frozen representations demonstrated their efficacy when the dataset is balanced, as evidenced in the IC-IT case.

This finding is substantiated by S2 File, which depict superior performance with frozen representation on the balanced dataset. However, the difference was not statistically significant (with a p-value ¿ 0.05) across most of the PLMs, rendering this observation as noteworthy, though not decisive.

The observed phenomenon intriguingly suggests a potential connection between dataset balance and the concepts of frozen and fine-tuned representations. Rather than treating these concepts as mutually exclusive, our study proposes that different tasks may warrant exploration of varying combinations of these methodologies, indicating the necessity for a more nuanced approach in their application.

#### Size of PLMs and performance

Our findings challenge the prevailing notion that the performance of PLMs invariably scales in direct proportion to their size. Interestingly, we did not identify a clear linear correlation between the dimensionality of a PLM and its ensuing performance. As a case in point, ESM-1b, with its 650 million parameters, consistently outperformed ESM-2 15B, which boasts 15 billion parameters, even when dealing with frozen representations (refer to S1 File). This observation underscores the conclusion that the performance efficacy of a PLM does not hinge exclusively on its size. Instead, it is shaped by a more intricate interplay of factors, with architectural design playing a significant role.

### Comparative performance analysis of classifiers

Table 5 presents performance results grouped by various classifiers utilized for three distinct protein classification tasks: distinguishing IC from MP, differentiating IT from MP, and discerning IC from IT.

**Table 5.** Performance overview of classifiers across protein classification tasks. This table offers a comprehensive performance evaluation of each classifier across three distinct protein classification tasks: differentiating ion channels (IC) from membrane proteins (MP), distinguishing ion transporters (IT) from MPs, and discerning IC from IT. The results, captured via a 5-fold cross-validation approach, are represented as mean±standard deviation. An accompanying p-value quantifies the statistical significance of observed differences among the classifiers. The highest value achieved for each task and column is marked in bold, while the second highest value is underlined to facilitate a comparison between the top-performing models.

| Task | Classifier | MCC | Accuracy | Sensitivity | Specificity | P-value |
| --- | --- | --- | --- | --- | --- | --- |
| IC-MP | LR | 0.82±0.04 | 93.99±1.30 | 85.53±4.03 | 94.69±1.97 | 2.29e-14 |
|  | kNN | 0.68±0.05 | 87.52±1.71 | 82.96±4.62 | 82.13±2.68 |  |
|  | SVM | **0.84±0.04** | **94.51±1.13** | 85.76±3.69 | 95.66±1.71 |  |
|  | RF | 0.69±0.05 | 92.00±1.38 | 63.96±4.59 | **96.86±1.52** |  |
|  | FFNN | 0.83±0.04 | **94.10±1.19** | **86.66±3.93** | 94.66±1.82 |  |
|  | CNN | 0.83±0.05 | 93.96±1.93 | 85.07±5.63 | 95.40±3.84 |  |
| IT-MP | LR | 0.80±0.04 | **93.12±1.34** | 83.71±3.74 | 94.19±2.21 | 4.77e-11 |
|  | kNN | 0.69±0.05 | 88.54±1.76 | 80.58±4.21 | 85.93±2.56 |  |
|  | SVM | **0.81±0.04** | **93.17±1.21** | **84.28±4.47** | 94.62±1.96 |  |
|  | RF | 0.65±0.05 | 90.33±1.62 | 64.35±4.47 | 93.57±2.14 |  |
|  | FFNN | **0.81±0.04** | **93.19±1.41** | **84.61±4.04** | 94.03±2.43 |  |
|  | CNN | **0.81±0.04** | **93.70±1.15** | 82.66±4.80 | **95.23±2.14** |  |
| IC-IT | LR | 0.82±0.03 | 91.22±1.61 | 91.00±3.11 | 91.44±3.44 | 1.38e-17 |
|  | kNN | 0.74±0.06 | 86.44±3.22 | 89.83±4.33 | 83.56±5.56 |  |
|  | SVM | 0.85±0.04 | **92.28±1.67** | 91.67±3.61 | **93.00±3.56** |  |
|  | RF | 0.79±0.04 | 89.28±2.22 | 86.28±6.06 | 91.72±6.06 |  |
|  | FFNN | 0.84±0.04 | **92.06±2.17** | **92.11±3.56** | 92.11±3.94 |  |
|  | CNN | **0.86±0.03** | **92.67±1.67** | 91.61±3.17 | **93.78±3.39** |  |

Our comprehensive investigation across distinct protein classification tasks, employing various classifiers, revealed a number of compelling insights.

#### Prominence of SVM and CNN classifiers

Both the Support Vector Machine (SVM) and Convolutional Neural Network (CNN) classifiers consistently delivered superior performance across all tasks. These classifiers effectively navigate high-dimensional data and unravel complex patterns, contributing to their consistent performance. The CNN employs convolutional layers to identify local patterns in the representations and nonlinear relationships inherent in neural network layers, while the SVM excels at linear classification by distinguishing between classes efficiently by maximizing margins.

#### Comparison of simple and complex models

Interestingly, a comparison of simple models, such as Logistic Regression (LR), and complex ones, like CNNs, indicated comparable performance levels. This observation counters the prevalent assumption that increasing model complexity necessarily results in superior performance. The consistent trend across all tasks and evaluation metrics suggests that in predicting IC and IT from MP, simpler models may deliver effectiveness on par with their more complex counterparts.

#### Less effective classifiers

However, not all classifiers showcased this level of effectiveness. Classifiers such as the k-Nearest Neighbors (kNN) and Random Forest (RF) were identified as the least effective across these tasks and representations derived from PLMs. This finding suggests that these classifiers may not align well with the specific nature of these tasks or the representations provided by the PLMs.

#### Performance parallels among classifiers

Furthermore, our analysis disclosed an intriguing parallel in the performance metrics of LR and Feed-Forward Neural Networks (FFNN), and those of SVM and CNN. This pattern suggests that, despite inherent differences in their complexity and structure, these models can achieve similar results in these specific tasks.

#### Significance of classifier selection

Finally, the p-value analysis highlighted significant performance differences across the classifiers for all three tasks, emphasizing the crucial role of classifier selection in the outcomes of these prediction tasks. The observed variation implies that the effectiveness of a specific classifier may vary based on the unique characteristics of the task, underscoring the importance of thoughtful classifier selection.

### Effects of various experimental conditions

In this section, we delve deeper into our findings and their implications. We have conducted three distinct assessments to elucidate their impacts on the results and overall performance. The following subsections offer a comprehensive discussion on these critical areas of impact, namely, the implications of frozen vs fine-tuned representations, the influence of balanced vs imbalanced datasets, and the effects of half vs full precision floating-point computations.

#### Frozen vs fine-tuned PLM representations

Table 6 presents the impact of frozen and fine-tuned representations across the three tasks under consideration - IC-MP, IT-MP, and IC-IT. Additionally, Fig 2 underscores the performance, specifically focusing on the MCC metric across the three tasks. Note that a comprehensive analysis concerning the influence of frozen and fine-tuned representations is available in S1 File.

**Table 6.**
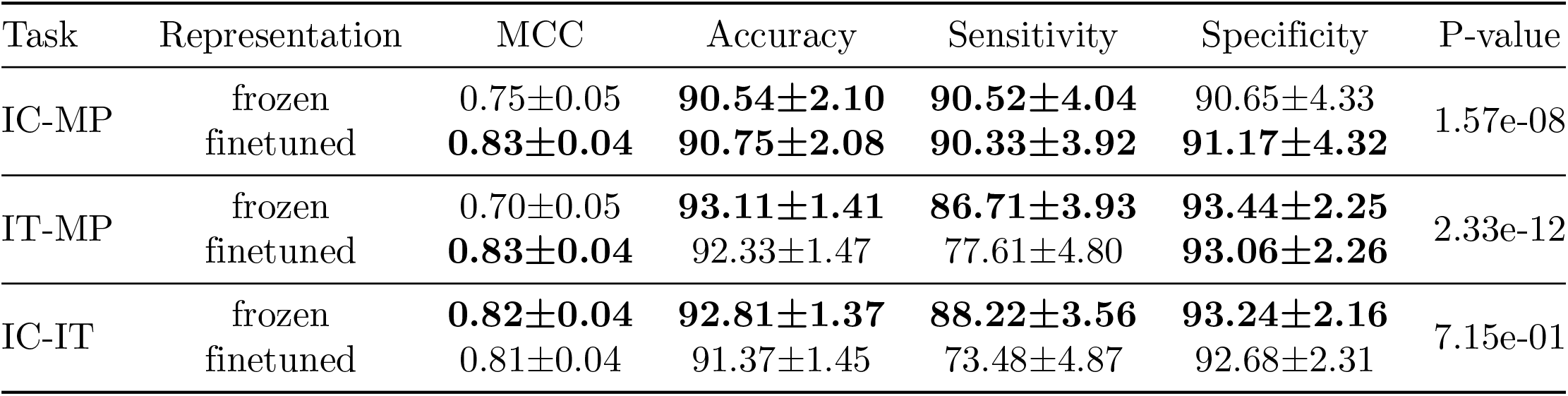
Comparison and evaluation of frozen and fine-tuned representations across diverse protein language models (PLMs). This table delineates the impact of utilizing both frozen and fine-tuned representations on three distinct tasks: differentiating Ion Channels (IC) from Membrane Proteins (MP), segregating Ion Transporters (IT) from MPs, and discriminating IC from IT, utilizing a range of PLMs. Four evaluation metrics have been computed using 5-fold cross-validation, presented as mean±standard deviation. The p-value is provided as a metric of the statistical significance of observed discrepancies among the models. Notably, the highest performance value for each task and each column is highlighted in boldface.

| Task | Representation | MCC | Accuracy | Sensitivity | Specificity | P-value |
| --- | --- | --- | --- | --- | --- | --- |
| IC-MP | frozen | 0.75±0.05 | **90.54±2.10** | **90.52±4.04** | 90.65±4.33 | 1.57e-08 |
|  | finetuned | **0.83±0.04** | **90.75±2.08** | **90.33±3.92** | **91.17±4.32** |  |
| IT-MP | frozen | 0.70±0.05 | **93.11±1.41** | **86.71±3.93** | **93.44±2.25** | 2.33e-12 |
|  | finetuned | **0.83±0.04** | 92.33±1.47 | 77.61±4.80 | **93.06±2.26** |  |
| IC-IT | frozen | **0.82±0.04** | **92.81±1.37** | **88.22±3.56** | **93.24±2.16** | 7.15e-01 |
|  | finetuned | 0.81±0.04 | 91.37±1.45 | 73.48±4.87 | 92.68±2.31 |  |

**Fig 2.**
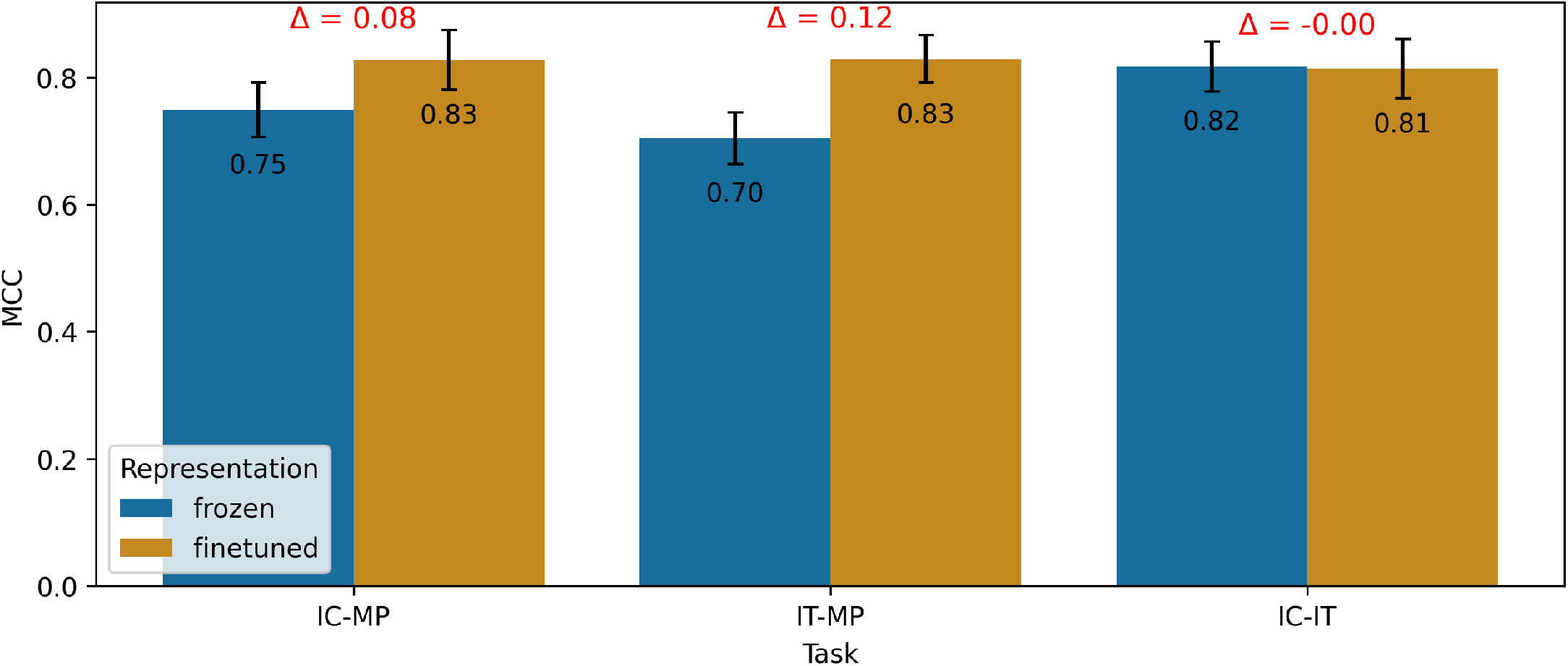
Graphical representation of the impact of frozen vs fine-tuned representations on various tasks across different Protein Language Models (PLMs). This figure elucidates the impact of employing frozen and fine-tuned representations across a range of Protein Language Models (PLMs) for three distinct tasks: differentiating Ion Channels (IC) from Membrane Proteins (MP), distinguishing Ion Transporters (IT) from MPs, and discriminating IC from IT. The results are portrayed using the mean Matthew’s Correlation Coefficient (MCC) values derived from 5-fold cross-validation. Each bar represents the mean MCC calculated across five cross-validation runs, while the error bars indicate the associated standard deviation. The symbol Δ is employed to denote the disparity between the corresponding pair of bars.

Our investigation has uncovered noteworthy disparities in the performance of fine-tuned and frozen representations across various tasks, underscored by their responses to task-specific conditions, dataset sizes, classifier choices, and the underlying PLM’s architecture.

##### Task-specific performance variations and the impact of dataset imbalances

On differentiating IC from MP and IT from MP, fine-tuned representations have consistently outperformed frozen ones. This pattern, however, becomes less clear-cut in the IC-IT task. Statistical analysis further supports this pattern, revealing substantial performance discrepancies between frozen and fine-tuned representations in the IC-MP and IT-MP tasks. However, the IC-IT task showed no significant difference.

This relative performance convergence in the IC-IT task can be attributed to the balanced nature of its dataset, contrasting with potential imbalances in the MP dataset. This highlights the role of dataset balance in performance trends and suggests that evaluation metrics may capture varying aspects of model performance, particularly under conditions of dataset imbalance.

A case in point is the sensitivity metric for the IT-MP task. Here, frozen representations notably outshine their fine-tuned counterparts, contrasting with the general trend of fine-tuned superiority. This demonstrates the sensitivity metric’s specific susceptibility to the effects of dataset imbalance. Whereas MCC metric, which accounts for all types of prediction errors, demonstrated equivalent performance for both representation types.

##### Influence of dataset size on performance

Our analysis points towards a significant influence of dataset size on the performance of fine-tuned representations. The larger, albeit imbalanced, MP dataset, comprising 3,413 training sequences, rendered richer fine-tuned representations compared to the balanced dataset of 280 sequences. Consequently, the benefits of fine-tuning appear more distinct with larger datasets, underscoring the potential of using extensive data resources to enhance fine-tuned PLM representation performance.

The observed pattern suggests that larger models, such as ProtT5 and ESM-2 15B—currently unexplored due to computational limitations—could potentially exhibit improved performance given the feasibility of fine-tuning.

##### Performance across different classifiers

A further probe into performance across all classifiers, as represented in S1 File, demonstrated the consistent outperformance of fine-tuned over frozen representations. This observation reinforces the role of fine-tuning as a potent strategy to optimize PLM effectiveness across varied classifier architectures.

##### Performance across diverse PLMs

Our findings, as showcased in S1 File, reveal that performance remains relatively stable between diverse PLM sizes when using frozen representations. However, ESM-1b, a larger model with 650M parameters, outperformed smaller-sized PLMs like ProtBERT with 420M parameters. This observation suggests that the size of the underlying PLM can exert influence on the performance of frozen representations.

#### Balanced vs imbalanced datasets

Table 7 and Fig 3 present the performance of the six PLMs when applied to either a balanced or imbalanced MP dataset. Our analysis suggests a profound effect of dataset balance on the performance of different representations across PLMs, classifiers, and tasks.

**Table 7.** Performance of Protein Language Models (PLMs) on Balanced vs Imbalanced Membrane Protein Datasets. This comprehensive evaluation examines the performance of various Protein Language Models (PLMs) on both balanced and imbalanced datasets of membrane proteins. The results, computed using 5-fold cross-validation, are represented as mean±standard deviation for the evaluation metrics. The p-value quantifies the statistical significance of observed differences amongst the classifiers. The highest values for each task and column are highlighted in bold. The PLMs are sorted based on their number of parameters.

| PLM | Dataset | MCC | Accuracy | Sensitivity | Specificity | P-value |
| --- | --- | --- | --- | --- | --- | --- |
| ProtBERT | balanced | 0.70±0.06 | 89.14±2.42 | **89.00±3.33** | 89.27±3.89 | 2.52e-01 |
|  | imbalanced | **0.74±0.04** | **98.48±0.06** | 84.52±3.52 | **99.58±0.10** |  |
| ProtBERT-BFD | balanced | 0.71±0.06 | 88.55±2.45 | **88.94±3.61** | 88.24±4.12 | 1.57e-02 |
|  | imbalanced | **0.77±0.03** | **97.98±0.19** | 78.79±5.02 | **99.56±0.08** |  |
| ESM-1b | balanced | 0.79±0.05 | 87.83±2.52 | **89.81±3.38** | 85.87±3.78 | 1.38e-04 |
|  | imbalanced | **0.87±0.02** | **96.92±0.17** | 67.83±5.25 | **99.17±0.25** |  |
| ESM-2 | balanced | 0.78±0.05 | 84.82±2.94 | **85.83±4.14** | 83.87±5.04 | 9.25e-03 |
|  | imbalanced | **0.83±0.03** | **96.92±0.23** | 66.71±5.44 | **99.40±0.12** |  |
| ProtT5 | balanced | **0.79±0.05** | 85.31±3.19 | **85.59±4.54** | 85.07±4.77 | 4.33e-01 |
|  | imbalanced | 0.75±0.04 | **97.25±0.08** | 69.50±5.06 | **99.50±0.17** |  |
| ESM-2.15B | balanced | **0.77±0.05** | 89.08±2.35 | **89.32±3.48** | 88.77±3.67 | 6.05e-01 |
|  | imbalanced | 0.73±0.03 | **96.83±0.17** | 67.92±5.92 | **99.17±0.08** |  |

**Fig 3.**
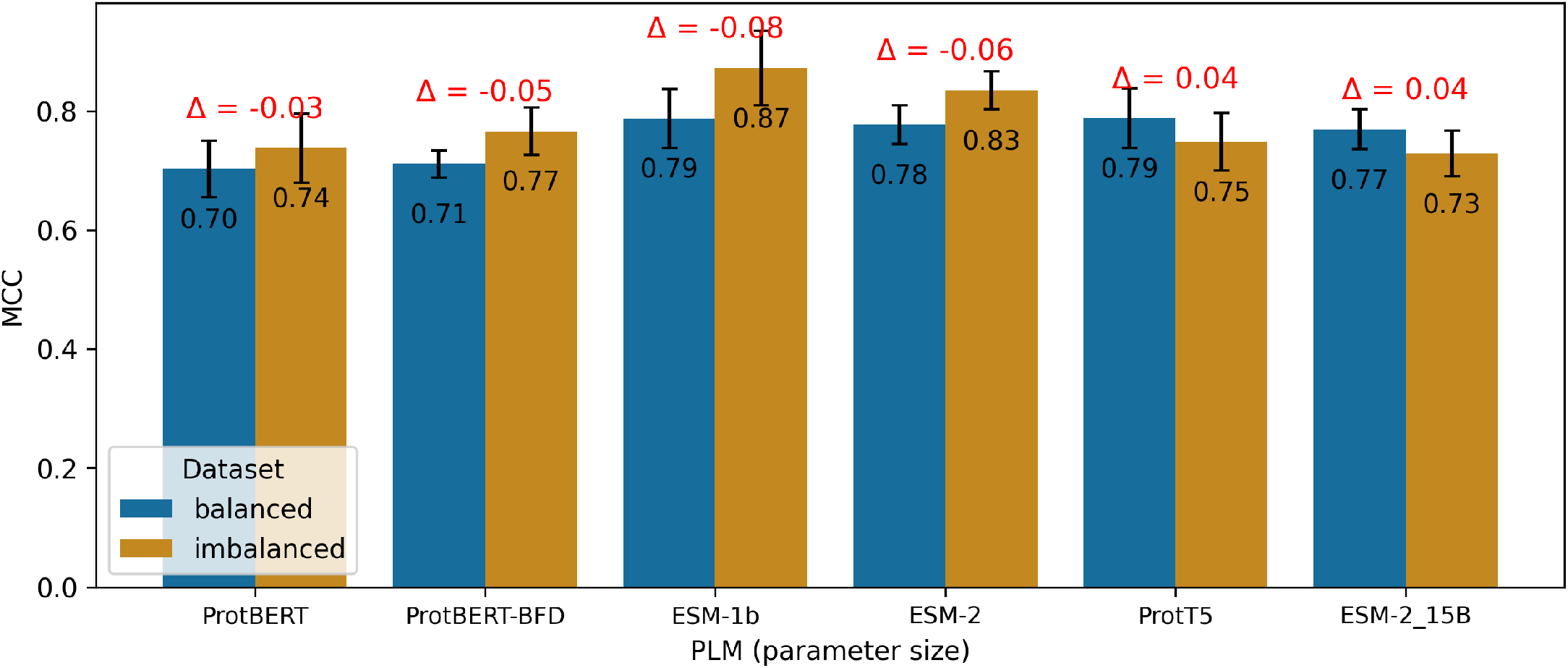
Evaluation of PLMs on balanced and imbalanced datasets of membrane proteins. This figure showcases a comprehensive evaluation of various protein language models (PLMs) on both balanced and imbalanced datasets of membrane proteins. The evaluation results are depicted as the mean Matthews Correlation Coefficient (MCC) calculated over five cross-validation runs, with error bars denoting the standard deviation. The symbol Δ indicates the difference between the corresponding pair of bars, providing insights into the performance disparities across the evaluated PLMs.

##### Performance across PLMs

Our results, as presented in Table 7 and Fig 3, indicate that representations from imbalanced datasets outperform those from balanced datasets across six PLMs, with the exception of ProtT5 and ESM-2 15B. This inconsistency may arise from the lack of fine-tuned representations for these specific PLMs. Given the feasibility of fine-tuning, we expect that these PLMs would align with the overall trend, affirming the performance advantage of imbalanced datasets.

However, the reported p-value in Table 7 suggests no significant difference between balanced and imbalanced datasets for ProtBERT, ProtT5, and ESM-2 15B PLMs. As ProtT5 and ESM-2 15B were not fine-tuned, the observed p-value primarily reflects the impact of dataset balance on the performance of frozen representations for these PLMs.

##### Task-specific performance variations

Evidence from results in S2 File indicates a superior performance of imbalanced datasets in the IC-MP and IT-MP tasks. These findings underscore the impact of dataset balance on model performance across these specific tasks.

##### Performance across different classifiers

The comparison of classifier performances presented in S2 File suggests that imbalanced datasets outshine balanced datasets across all classifiers, except for the RF classifier. This exception implies a particular sensitivity of the RF classifier to dataset balance, potentially explaining its performance divergence from the other classifiers.

##### Fine-tuned vs frozen representations

The performance patterns as seen in S2 File demonstrate that imbalanced datasets exhibit superior performance when employing fine-tuned representations across all fine-tuned PLMs. In contrast, balanced datasets perform better when using frozen representations, except for ProtBERT, where the p-value of 8.66e-02 indicates a statistically significant difference. These findings emphasize the significant impact of dataset balance on model performance, dependent on the choice of representation type (fine-tuned or frozen).

#### Half vs full precision floating point calculations

Table 8 and Fig 4 present the outcomes obtained from employing half and full precision floating-point calculations across the classifiers. Our analysis explores the influence of numerical precision—specifically half versus full precision floating-point calculations—on the performance of different tasks, classifiers, and PLMs.

**Table 8.** Performance of half vs full precision floating-point across six classifiers. This table provides an overview of the performance of each classifier using half and full precision floating-point calculations. The results are presented using evaluation metrics on the 5-fold cross-validation, with the mean and standard deviation shown. The value indicates the statistical significance of the observed differences among the classifiers. The highest value for each task and each column is highlighted in bold.

| Classifier | Precision | MCC | Accuracy | Sensitivity | Specificity | P-value |
| --- | --- | --- | --- | --- | --- | --- |
| LR | half | **0.82±0.04** | **93.56±1.62** | **85.54±5.08** | 94.99±3.04 | 9.69e-01 |
|  | full | 0.81±0.04 | **93.62±1.53** | **85.32±4.58** | **95.02±3.10** |  |
| kNN | half | 0.69±0.05 | **93.20±1.50** | **86.95±3.90** | **93.78±2.56** | 9.01e-01 |
|  | full | **0.70±0.05** | **93.43±1.45** | **86.92±3.90** | **93.99±2.44** |  |
| SVM | half | **0.83±0.04** | 92.93±1.42 | **85.91±3.80** | 93.65±2.44 | 9.22e-01 |
|  | full | **0.83±0.04** | **93.22±1.35** | **85.88±3.68** | **94.00±2.29** |  |
| RF | half | 0.69±0.05 | **90.81±1.73** | **70.20±5.09** | **94.31±2.76** | 9.64e-01 |
|  | full | **0.70±0.05** | **90.77±1.58** | 67.29±4.63 | **94.68±2.61** |  |
| FFNN | half | **0.82±0.04** | **93.40±1.28** | **86.41±3.98** | **94.59±2.24** | 9.27e-01 |
|  | full | **0.82±0.04** | **93.63±1.26** | **86.30±3.99** | **94.81±2.13** |  |
| CNN | half | **0.83±0.04** | **87.85±2.04** | **83.29±4.37** | **84.35±3.19** | 8.09e-01 |
|  | full | **0.83±0.04** | **87.60±2.03** | **83.46±4.42** | **83.60±3.22** |  |

**Fig 4.**
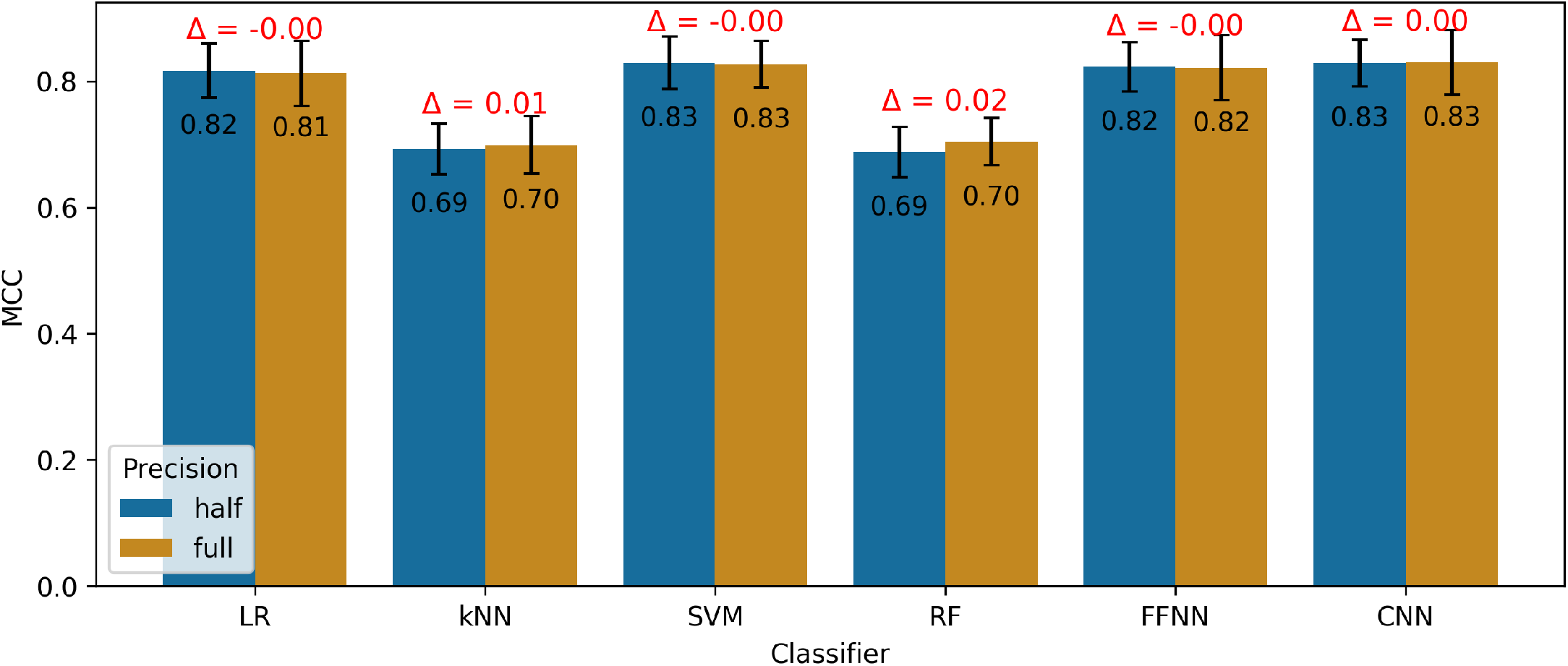
Half vs full precision evaluation across classifiers. This evaluation compares the performance of different protein language models (PLMs) using both half and full precision floating-point calculations. The results are presented as the mean Matthews Correlation Coefficient (MCC) calculated across five cross-validation runs, with error bars indicating the standard deviation. The symbol Δ represents the difference between the corresponding pair of bars, providing insights into the impact of numerical precision on classifier performance.

##### Performance across different classifiers

As evidenced by the results presented in Table 8 and Fig 4, the performance remains consistent across all classifiers, irrespective of whether half or full precision floating-point calculations are employed. This suggests that the level of numerical precision does not significantly affect classifier performance in the evaluated tasks.

##### Task-specific performance variations

Performance consistency extends to specific tasks as well. As shown in S3 File, the IC-MP, IT-MP, and IC-IT tasks exhibit comparable performance levels regardless of the employed floating-point precision. These findings reinforce the notion that the numerical precision choice for the floating-point calculations does not materially affect model performance across these tasks.

##### Performance across PLMs

The performance comparison among the six PLMs, as displayed in S3 File, reveals minor performance variations when using both half and full precision floating-point calculations. This observation implies that the selection of floating-point precision has minimal impact on the performance of the evaluated PLMs.

##### Influence on evaluation metrics and statistical significance

An overarching analysis of evaluation metrics and p-values reveals no statistically significant differences between the usage of half and full precision floating-point calculations across varied tasks, classifiers, and PLMs. These findings underscore that the choice of floating-point precision does not exert a considerable influence on the outcomes of the prediction tasks assessed in this study.

#### Visualization of representations: insights and implications

The UMAP projection matrix of representations derived from the ESM-1b PLM, presented in Fig 5, provides a compelling visualization of both frozen and fine-tuned representations for balanced and imbalanced datasets within the context of the IC-MP task on the training set. It is crucial to note that the representation shown for the balanced dataset is randomly selected from one of the ten available balanced datasets.

**Fig 5.**
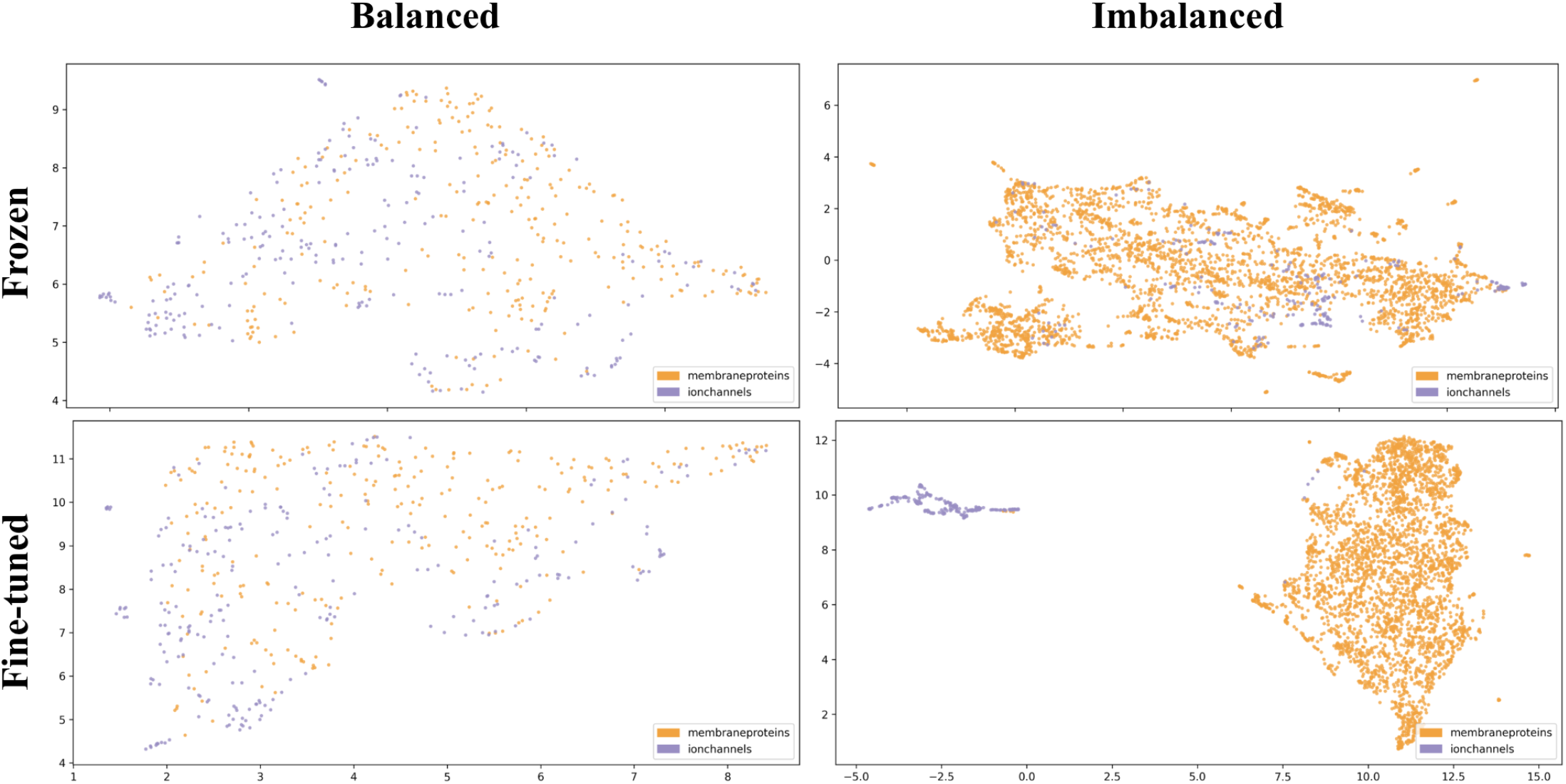
UMAP projection of representations from top PLM for ion channel discrimination. The figure showcases a UMAP projection of representations derived from ESM-1b, the highest-performing Protein Language Model (PLM) in the task of discriminating ion channels (IC) from membrane proteins (MP). The representations are visualized in four variations: frozen and fine-tuned representation types, along with balanced and imbalanced datasets. In the visualization, membrane protein representations are depicted in yellow, while ion channel protein representations are depicted in blue.

#### Fine-tuned representations in imbalanced dataset

The Fig 5 visualization underscores the distinct clusters and patterns within the fine-tuned representations for the imbalanced dataset. The evident separation between ion channels and membrane proteins signifies the highly discriminative capability of fine-tuned representations, demonstrating their efficacy in this task. This insight underscores the prowess of fine-tuned representations in capturing the unique and distinguishable characteristics of ion channels, fostering precise classification and analysis.

#### Frozen representations in imbalanced dataset

Notably, the visualization also indicates that the next best level of clarity is achieved using frozen representations with the imbalanced dataset. This suggests that the imbalanced dataset, enriched with a broader spectrum of other membrane proteins, enhances the performance of the frozen representations. This may be due to the diversity and complexity of the other membrane proteins, requiring a larger dataset for effective representation and discrimination. Hence, this highlights the advantage of employing imbalanced datasets with frozen representations for capturing the intricacies of diverse membrane protein structures.

#### Impact of undersampling on classification task

Our results accentuate the potential adverse consequences of undersampling the dataset on the classification task performance. Undersampling, which reduces the dataset size, can impair the model’s ability to classify proteins accurately, underscoring the need for a sufficiently large dataset to ensure effective protein classification. A substantial dataset ensures the model’s exposure to diverse and representative examples, facilitating the learning of robust, discriminative patterns that generalize well to unseen data. Consequently, securing a substantial dataset is of paramount importance for achieving optimal performance in protein classification tasks.

#### Implications for balanced dataset representations

Examining the visualization of frozen and fine-tuned representations with balanced datasets, we find a lack of clear patterns. This signifies a less distinct characterization of ion channels compared to other membrane proteins, suggesting these representations may not effectively differentiate ion channels from other membrane proteins. This lack of clear patterns implies that the representations derived from balanced datasets may fail to capture unique features or discriminative information vital for robust ion channel classification. Hence, alternative representation strategies or dataset balancing techniques may warrant consideration to enhance model effectiveness.

#### Comprehensive visualization of PLMs

The representation visualizations for all six PLMs, including both frozen and fine-tuned representations for the IC-MP, IT-MP, and IC-IT tasks, are provided in S4 File. As shown, these visualizations offer a holistic view of the performance and discriminative abilities of various PLMs and representations for these tasks. These comprehensive visualizations allow for an in-depth understanding of how different PLMs capture the characteristics and separability of ion channels and other membrane proteins, illuminating their respective strengths and weaknesses.

### Overview of top cross-validation results

The top results obtained from the 5-fold cross-validation (CV) for each task are detailed in Table 9. Results are stratified by classifier and presented in the CV column, showing the mean and standard deviation over the five folds. While independent test set results are provided for comparative purposes, they do not contribute to the selection of the best model, ensuring a robust and unbiased evaluation of classifier performance.

**Table 9.** Top 5-fold CV results for each task and classifier, along with independent test set results. This table presents the best 5-fold cross-validation (CV) results for each task and classifier, as well as the corresponding results on the independent test set for comparison purposes. The tasks include discriminating ion channels (IC) from other membrane proteins (MP), ion transporters (IT) from MP, and IC against IT. The table displays the mean and standard deviation of the 5-fold CV results for each metric. The results for the IC-MP and IT-MP tasks are obtained from imbalanced datasets, while the dataset for the IC-IT task remains balanced. The best values for each task are shown in bold, and the second-best values are underlined. It is important to note that the independent test set results are provided solely for evaluating the models based on the CV results and not for selecting the best model, as the best models are chosen based on the CV results.

| Task | Representation | Representer | Dataset | Classifier | MCC |  |
| --- | --- | --- | --- | --- | --- | --- |
|  |  |  |  |  | CV | Independent |
| IC-MP | finetuned | ESM-1b | Imbalanced | SVM | <u>0.99±0.01</u> | **0.85** |
|  |  |  |  | RF | <u>0.98±0.01</u> | <u>0.84</u> |
|  |  |  |  | kNN | <u>0.99±0.01</u> | <u>0.83</u> |
|  |  |  |  | LR | **1.00±0.00** | **0.85** |
|  |  |  |  | FFNN | **1.00±0.01** | **0.85** |
|  |  |  |  | CNN | <u>0.99±0.01</u> | **0.85** |
| IT-MP | finetuned | ESM-1b | Imbalanced | SVM | **1.00±0.00** | 0.68 |
|  |  |  |  | RF | <u>0.99±0.01</u> | 0.67 |
|  |  |  |  | kNN | <u>0.99±0.01</u> | **0.70** |
|  |  |  |  | LR | **1.00±0.00** | <u>0.69</u> |
|  |  |  |  | FFNN | **1.00±0.01** | <u>0.67</u> |
|  |  |  |  | CNN | <u>0.99±0.01</u> | <u>0.69</u> |
| IC-IT | frozen | ESM-2_15B | Balanced | SVM | <u>0.88±0.03</u> | **0.88** |
|  | finetuned | ESM-1b |  | RF | <u>0.84±0.03</u> | 0.79 |
|  | frozen | ProtT5 |  | kNN | <u>0.81±0.03</u> | 0.75 |
|  | finetuned | ESM-1b |  | LR | <u>0.88±0.05</u> | 0.79 |
|  | frozen | ESM-2 |  | FFNN | <u>0.88±0.05</u> | 0.74 |
|  | finetuned | ESM-2 |  | CNN | **0.89±0.03** | <u>0.87</u> |

#### Superior performance of ESM-1b PLM in IC-MP and IT-MP tasks

As outlined in Table 9, the ESM-1b PLM, combined with fine-tuned representations and an imbalanced dataset, exhibits superior performance in the IC-MP and IT-MP tasks. The LR and FFNN classifiers, in particular, achieve a perfect MCC of 1.00, indicating flawless prediction on 5-fold CV. Other classifiers also present highly competitive results, with MCC values reaching 0.99, thereby emphasizing the exceptional efficacy of the ESM-1b PLM with fine-tuning and an imbalanced dataset.

#### Results from multiple PLMs in IC-IT task

The IC-IT task, employing a balanced dataset, sees a range of PLMs delivering notable results. The top-performing classifier, CNN, leverages the ESM-2 PLM with fine-tuned representations, yielding an impressive MCC of 0.89. Notably, larger PLMs like ProtT5 and ESM-2 15B produce comparable results to their smaller counterparts such as ESM-1b and ESM-2. This suggests that the size of the PLM does not necessarily influence performance enhancement for the IC-IT task.

#### Comparative performance of classifiers for IC-IT task

While the CNN classifier utilizing the ESM-2 PLM’s fine-tuned representations achieves the top result for the IC-IT task, other classifiers also demonstrate comparable performances. The SVM classifier with frozen representations from ESM-2 15B, the LR classifier with fine-tuned representations from ESM-1b, and the FFNN classifier with frozen representations from ESM-2 deliver similar results to the CNN classifier. This suggests that a diverse set of classifiers can deliver equivalent performance levels, depending on the selected PLM and representation type.

#### Comprehensive analysis of results

A detailed examination of the results for each task - IC-MP, IT-MP, and IC-IT - is provided in S5 File. Here, the evaluation metrics are delineated in detail across various tables for each task. This thorough breakdown offers an exhaustive and nuanced understanding of the performance of the employed models, classifiers, and representations. Delving into the evaluation metrics’ specifics enables readers to gain deeper insights into the results, providing valuable information for future research in the prediction of ion channels and ion transporters from other membrane proteins.

### Performance comparison with state-of-the-art projects

A detailed comparison of TooT-PLM-ionCT’s performance against state-of-the-art projects is provided in Table 10 and Fig 6 for the IC-MP, IT-MP, and IC-IT tasks. This analysis includes established methodologies such as DeepIon [35], MFPS CNN [22], and TooT-BERT-C [23], providing a comprehensive assessment of TooT-PLM-ionCT’s relative performance.

**Table 10.** Comparative performance of TooT-PLM-ionCT with state-of-the-art. This table presents a comparative analysis of the performance of TooT-PLM-ionCT, which consists of three separate systems with distinct architectures and trained on different datasets, and the state-of-the-art methods on the independent test set. These systems precisely classify membrane proteins (MP), ion channels (IC), and ion transporters (IT). Instances where corresponding studies and tools do not report on ion channel and ion transporter classification against each other are denoted by a “-” symbol. Performance indicators, such as accuracy and Matthews Correlation Coefficient (MCC), are highlighted: boldface for the highest values and underlined for the second-highest. This table underscores the distinct and task-specific nature of the TooT-PLM-ionCT systems.

| Task | Project | Encoder | Classifier | Accuracy | MCC |
| --- | --- | --- | --- | --- | --- |
| IC-MP | DeepIon [35] | PSSM | CNN | 86.53 | 0.37 |
|  | MFPS_CNN [22] | PSSM | CNN | <u>94.60</u> | <u>0.62</u> |
|  | TooT-BERT-C [23] | ProtBERT-BFD | LR | **98.24** | **0.85** |
|  | **TooT-PLM-ionCT** | ESM-1b | LR | **98.24** | **0.85** |
| IT-MP | DeepIon [35] | PSSM | CNN | 83.78 | 0.37 |
|  | MFPS_CNN [22] | PSSM | CNN | 93.30 | 0.59 |
|  | TooT-BERT-C [23] | ProtBERT-BFD | LR | <u>95.43</u> | <u>0.64</u> |
|  | **TooT-PLM-ionCT** | ESM-1b | LR | **95.98** | **0.69** |
| IC-IT | DeepIon [35] | - | - | - | - |
|  | MFPS_CNN [22] | - | - | - | - |
|  | TooT-BERT-C [23] | ProtBERT-BFD | LR | <u>85.38</u> | <u>0.71</u> |
|  | **TooT-PLM-ionCT** | ESM-2 | CNN | **93.07** | **0.87** |

**Fig 6.**
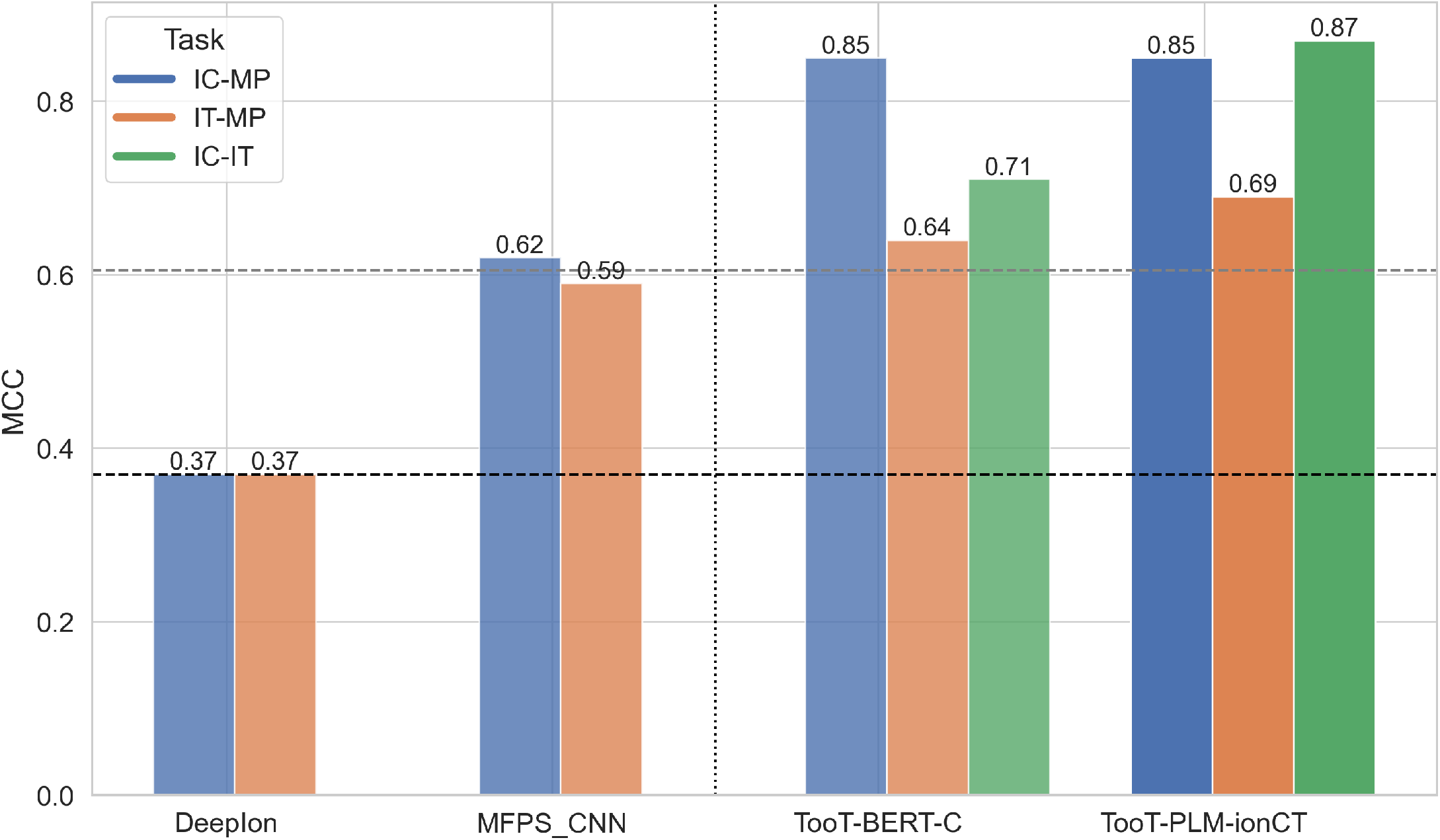
Comparative performance with state-of-the-art. This figure presents the comparative performance of TooT-PLM-ionCT on the independent test set, showcasing the classification results for membrane proteins (MP), ion channels (IC), and ion transporters (IT). The absence of bars indicates studies that focused on classifying ion channels and ion transporters against membrane proteins, rather than against each other, resulting in no available results from either publications or tools. The horizontal dashed lines represent two baselines, while the vertical dashed line distinguishes between traditional and PLM-based representations.

As shown in Table 10 and Fig 6, TooT-PLM-ionCT outperforms its counterparts in the IT-MP and IC-IT tasks. However, in the IC-MP task, its performance aligns closely with TooT-BERT-C. These results underscore the capability of TooT-PLM-ionCT to accurately predict ion channels and ion transporters from other membrane proteins, demonstrating its superiority or competitive performance.

It’s worth noting that DeepIon [35] and MFPS CNN [22] do not report specific results for the IC-IT task, as they focus predominantly on differentiating ion channels and ion transporters from other membrane proteins. This further underscores the unique contribution of our study in exploring the IC-IT task and offering crucial insights into the categorization of ion channels and ion transporters from other membrane proteins.

#### Model selection process

The model selection was driven by the top-performing models in our experiments, as detailed in Table 9. In instances where multiple classifiers achieved the same MCC, we favored the simpler and more straightforward classifier for the IC-MP and IT-MP tasks. However, for the IC-IT task, despite the SVM classifier’s marginally better performance on the independent test set, the CNN classifier was selected based on superior CV results. This decision balanced the need for performance with model simplicity, while considering the unique demands and constraints of each task.

## Conclusion

In this study, we introduced TooT-PLM-ionCT, a comprehensive framework consisting of three distinct systems, each specifically developed for a separate task: distinguishing ion channels (IC) from other membrane proteins (MP), ion transporters (IT) from MP, and IC from IT. Each system has its own architecture and was trained on a different dataset, underlining their task-specific nature.

The six Protein Language Models (PLMs) utilized were ProtBERT, ProtBERT-BFD, and ProtT5 from the ProtTrans project, along with ESM-1b, ESM-2 (650M parameters), and ESM-2 (15B parameters) from the ESM project. These were coupled with an array of both traditional classifiers (Logistic Regression, kNN, Random Forest (RF), SVM, and Feed-Forward Neural Network) and a deep learning (Convolutional Neural Network) classifier.

In our quest to understand PLM performance, we examined the impacts of dataset balance, the difference between frozen and fine-tuned representations, and the performance variance between half-precision and full-precision floating-point calculations. Our key findings from these analyses are as follows:

- **PLM Performance:** ESM-1b PLM outshone its peers in most metrics and tasks, with the exception of distinguishing IC from IT, where it shared the top spot with ESM-2 15B. The second-best performing model, however, varied with the task at hand. ESM-2 proved effective in differentiating IC from MP and IT from MP, while ProtT5 excelled in IC-IT classification. The substantial variation in p-values of statistical analysis across all PLMs further emphasized ESM-1b’s formidable performance.
- **Dataset Balance:** Our study found that imbalanced datasets outperformed balanced datasets across most PLMs, except for ProtT5 and ESM-2 15B, where we saw inconsistency due to the absence of fine-tuned representations. Additionally, a comparison of classifier performance revealed that imbalanced datasets outperformed balanced datasets across all classifiers. The sole exception was the RF classifier, which exhibited a heightened sensitivity to balanced datasets and therefore yielded superior results with them.
- **Fine-Tuned Representations:** Fine-tuned representations consistently performed better than frozen ones for differentiating IC from MP and IT from MP, while for the IC-IT task, the performance was equivocal. The size of the dataset appeared to significantly influence the performance of fine-tuned representations. Thus, larger datasets, despite their imbalanced nature, seemed to benefit more from fine-tuning.
- **Floating-Point Precision:** Our study found negligible performance variations between half and full precision floating-point calculations across tasks, classifiers, and PLMs. This suggests that the numerical precision choice does not considerably impact the performance in the prediction tasks examined in this study.
- **Impact of Undersampling:** Results highlighted the potential detrimental effects of undersampling, emphasizing the need for larger, more representative datasets for accurate protein classification.
- **Comparison of PLM Sizes:** Our analysis showed an intriguing pattern where a 650M-parameter PLM exhibited comparable performance to a 15B-parameter PLM and surpassed a 450M-parameter model in terms of frozen representation.
- **Computational Cost vs Improvement:** The improvement in performance for the IC-IT task justified the associated computational cost, a contrast to the IC-MP and IT-MP tasks where the benefit did not outweigh the cost.

In our future endeavors, we plan to further probe the distinct nature of each TooT-PLM-ionCT system, augmenting the representations produced by PLMs with additional sources of knowledge. We also aim to test more sophisticated sequence representation techniques and apply our approach on larger and more diverse protein datasets. These advancements will enable us to both validate the robustness of our task-specific systems and broaden their range of applicability. Our study has laid a solid foundation for such future exploration, bringing us a step closer to a more nuanced understanding of membrane proteins.

## Supporting information

**S1 File. Frozen vs Fine-tuned Representations.**

**S2 File. Balanced vs Imbalanced Datasets.**

**S3 File. Half vs Full Precision Floating Point Calculations.**

**S4 File. Protein visualization.**

**S5 File. Detailed five-fold cross-validation results.**

---

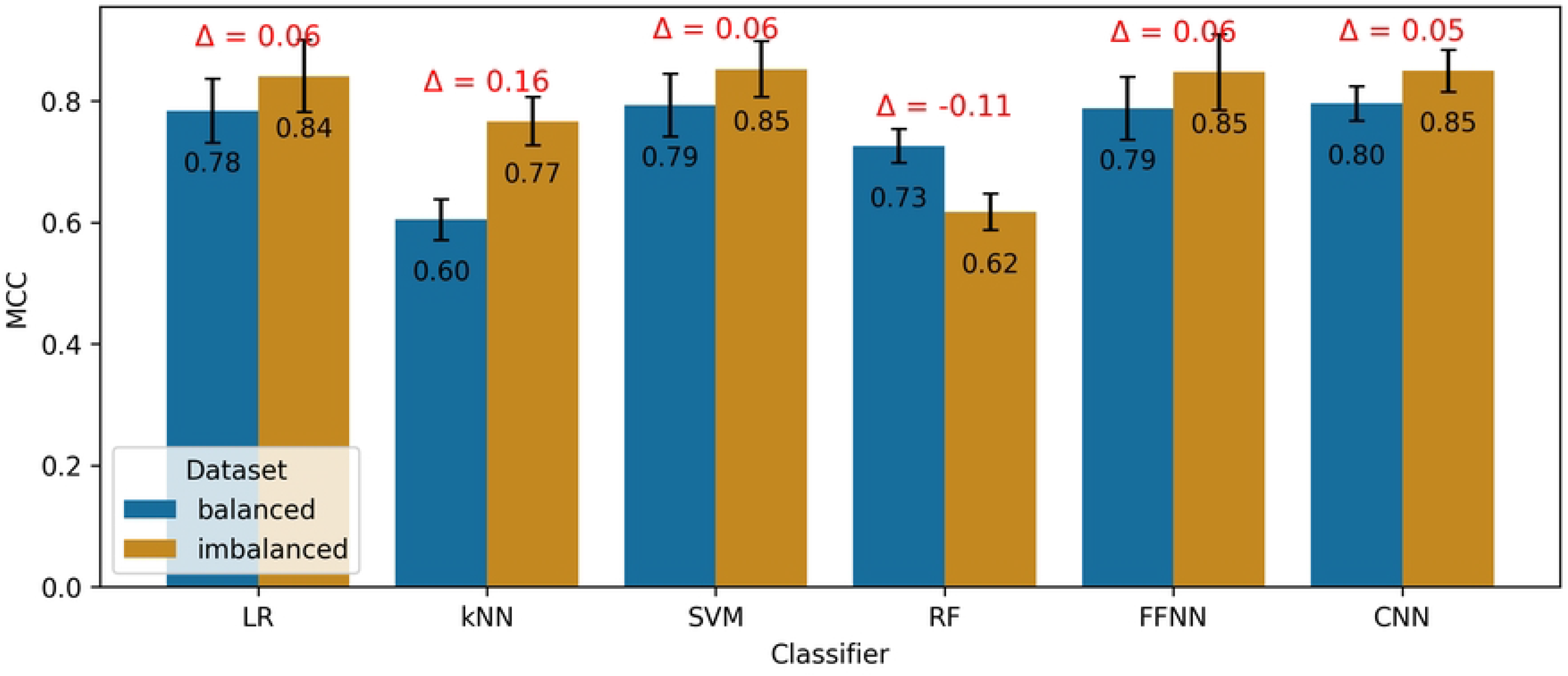

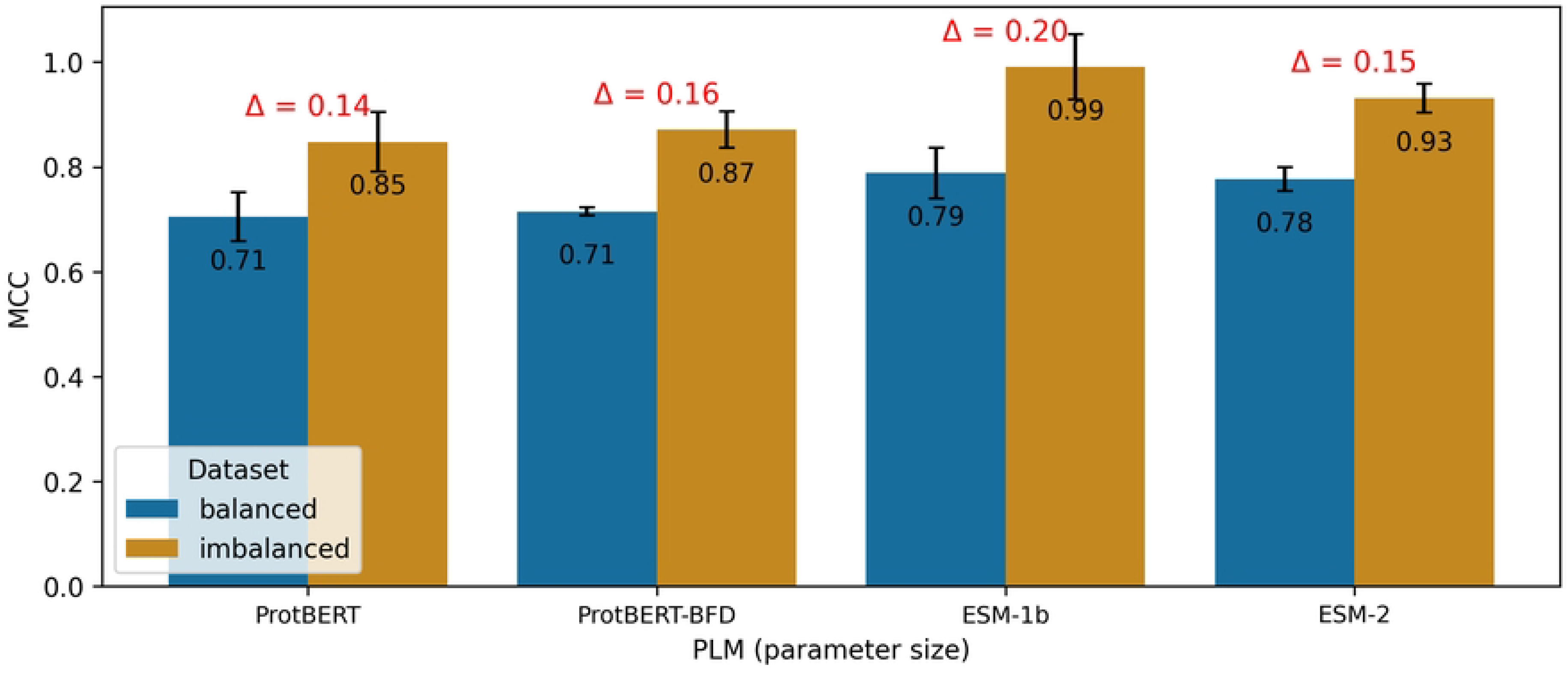

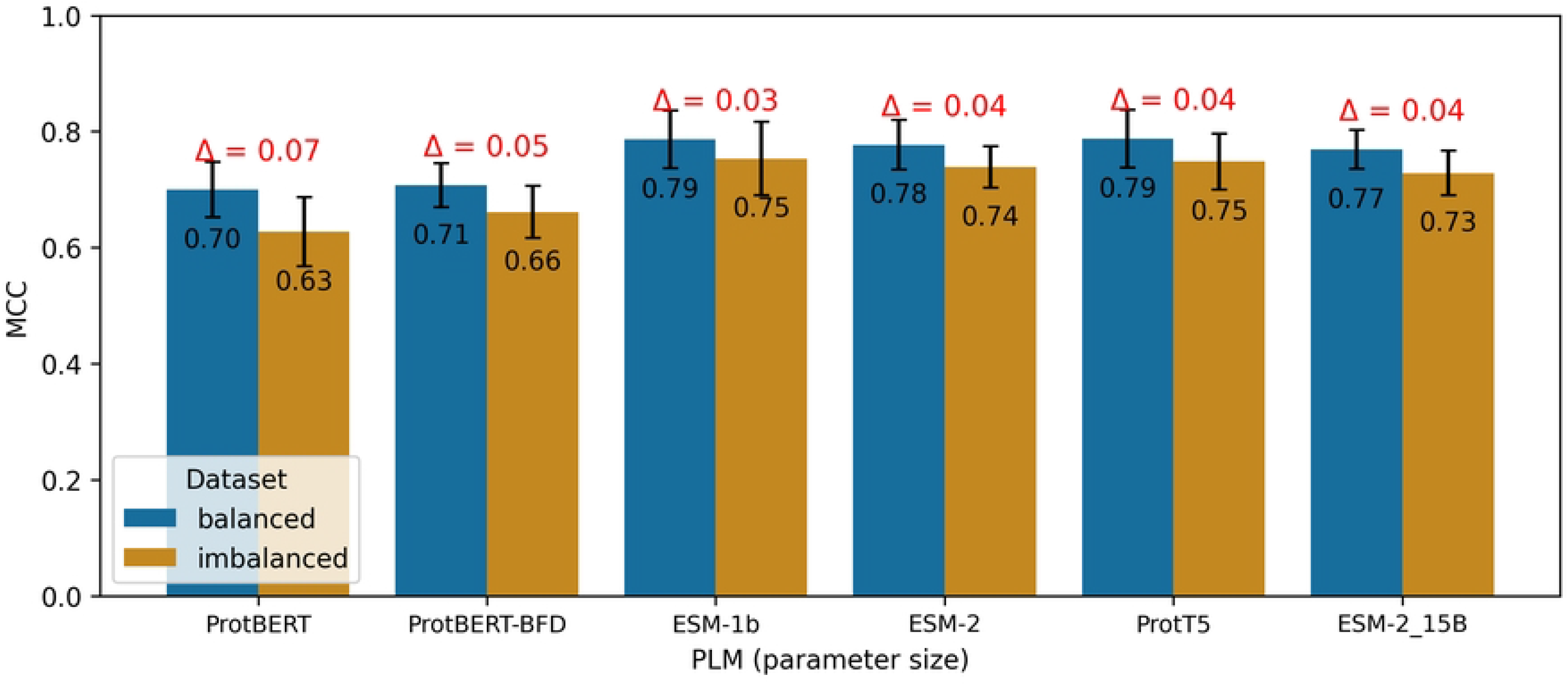

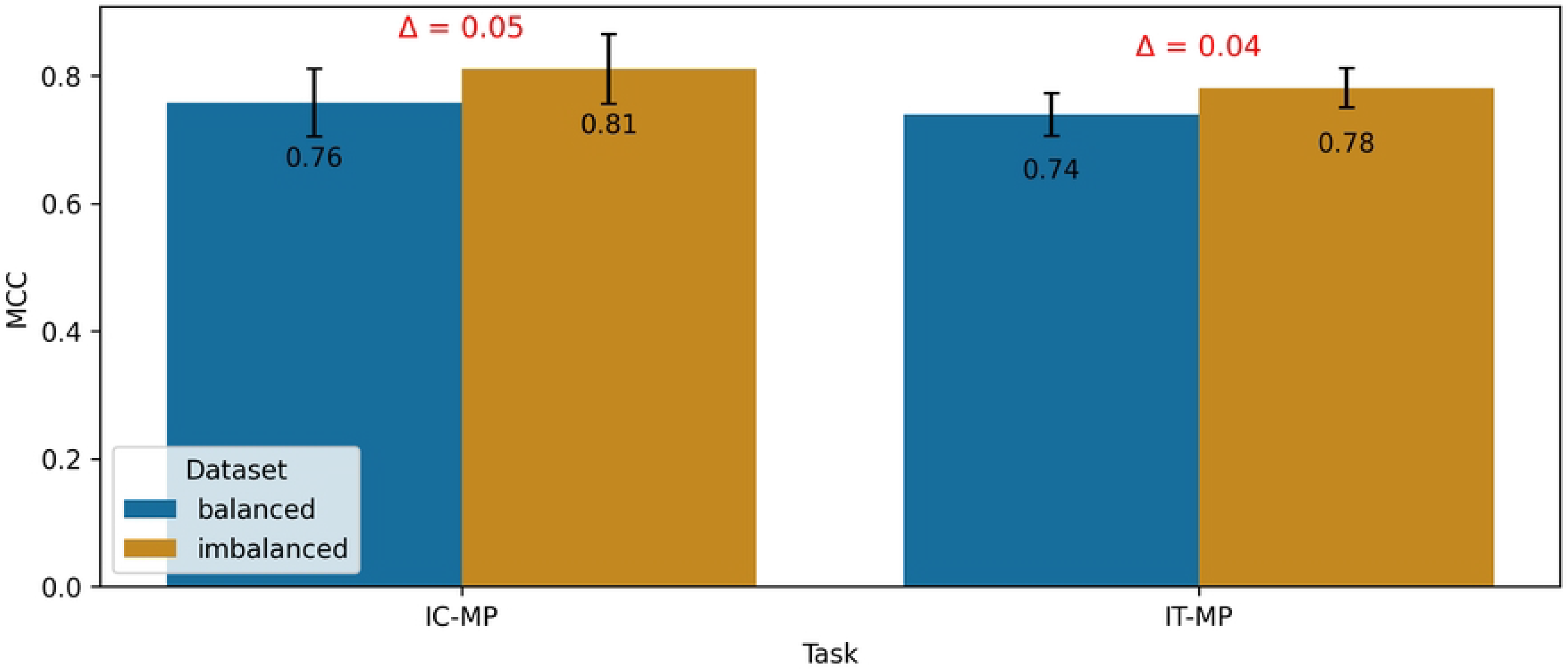

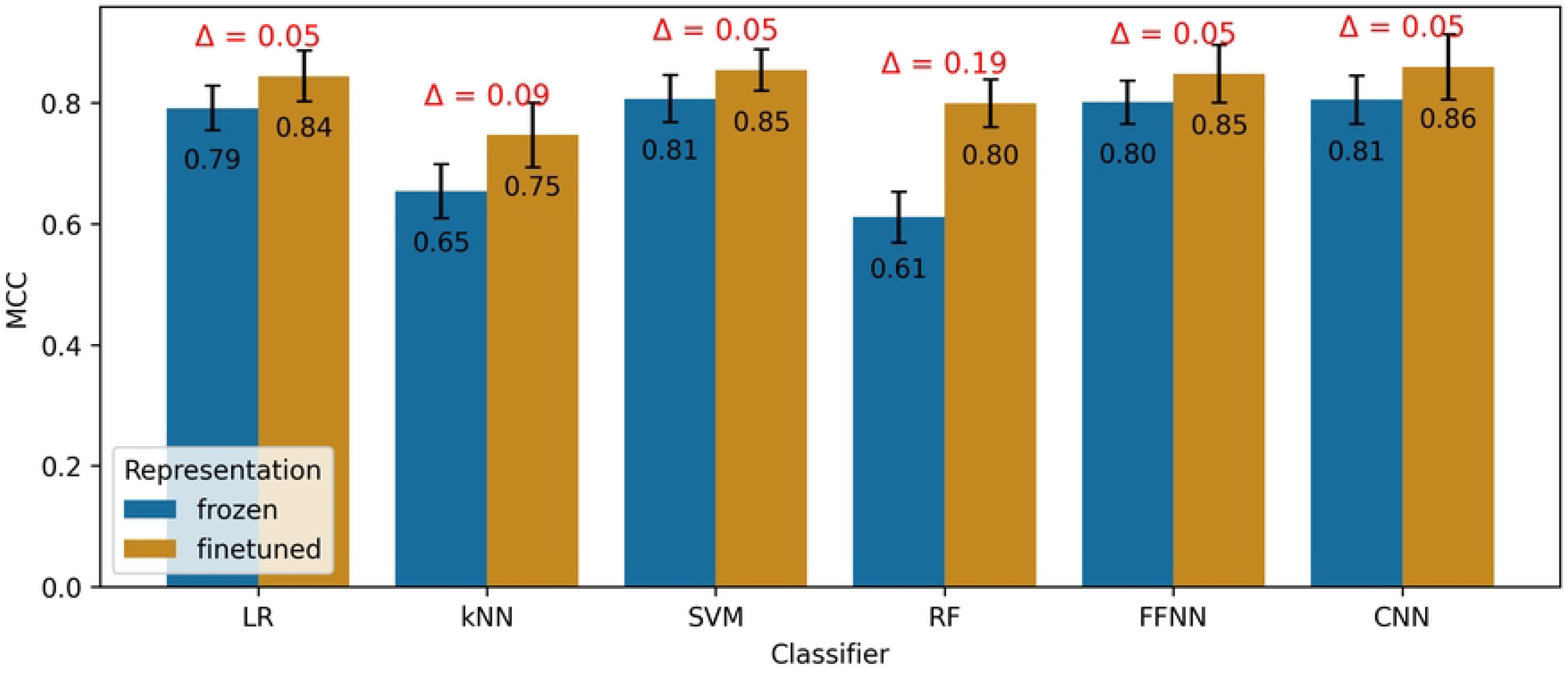

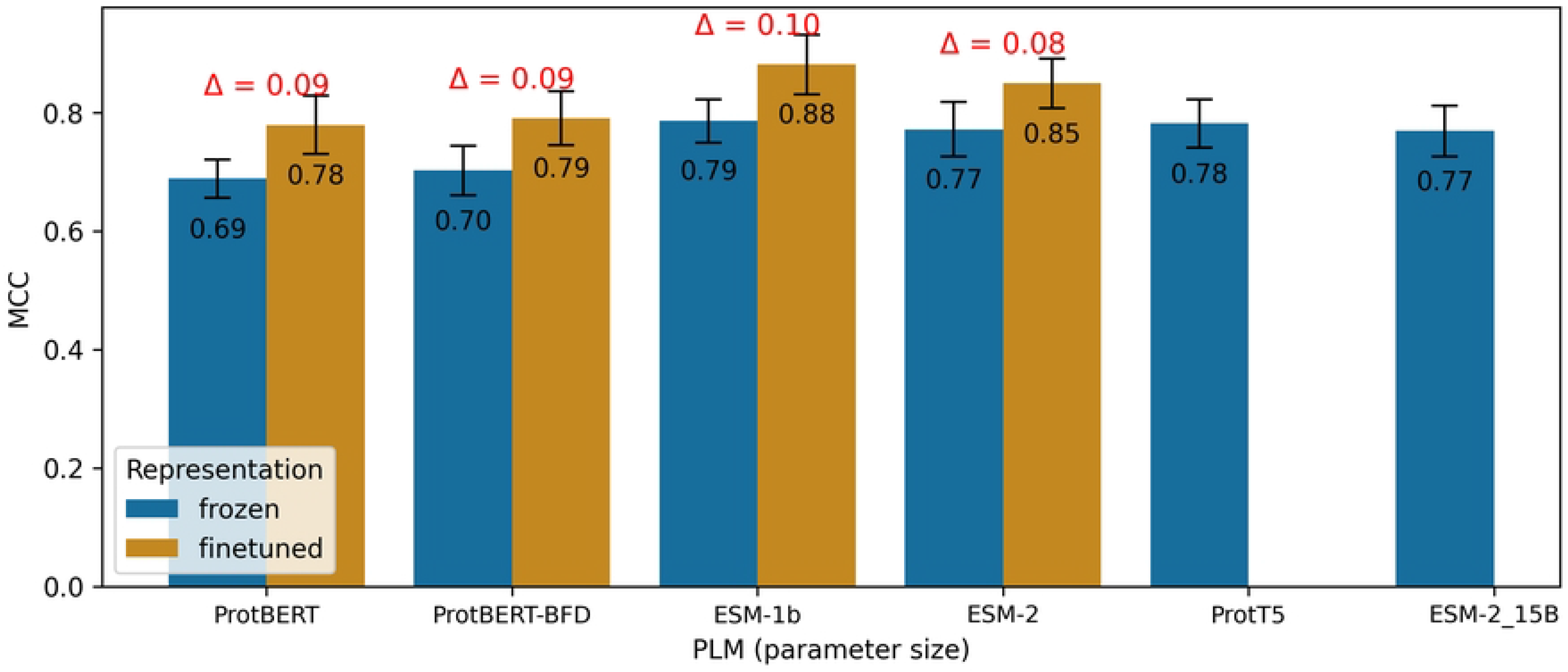

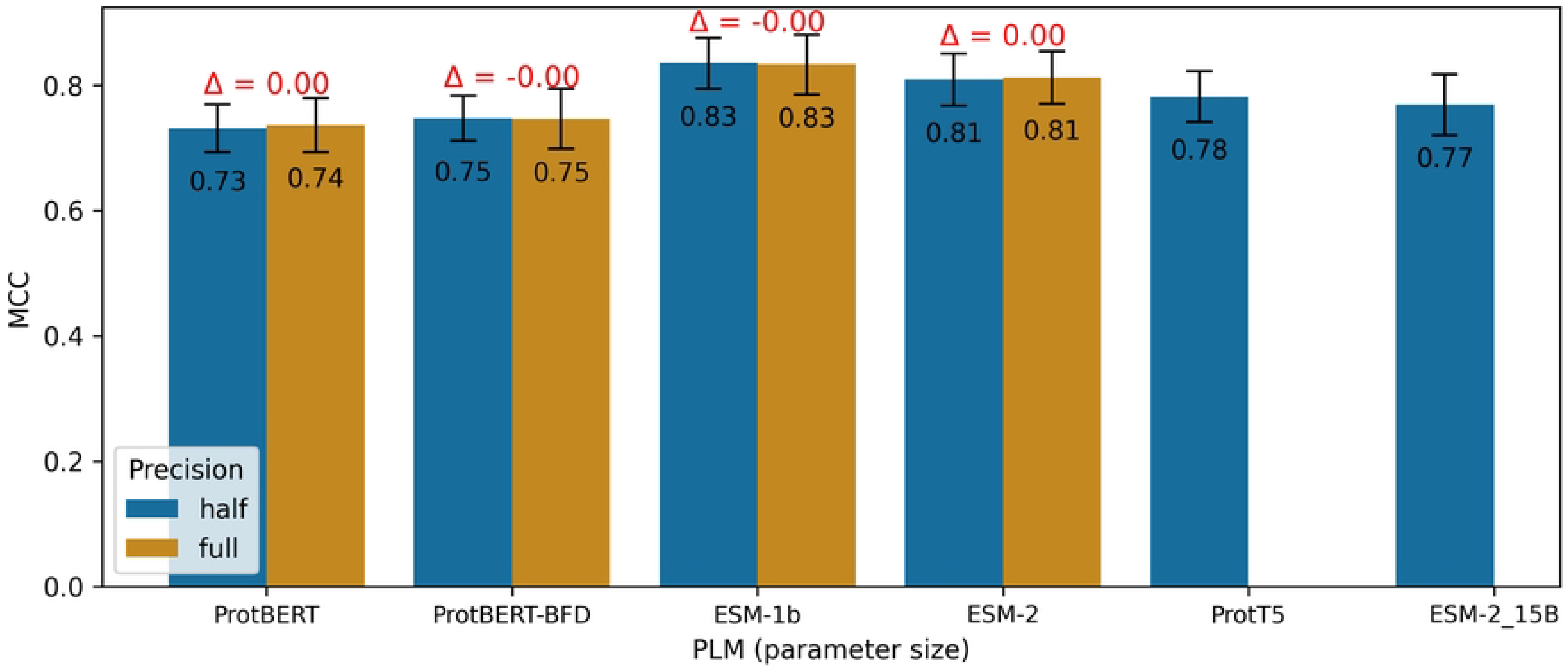

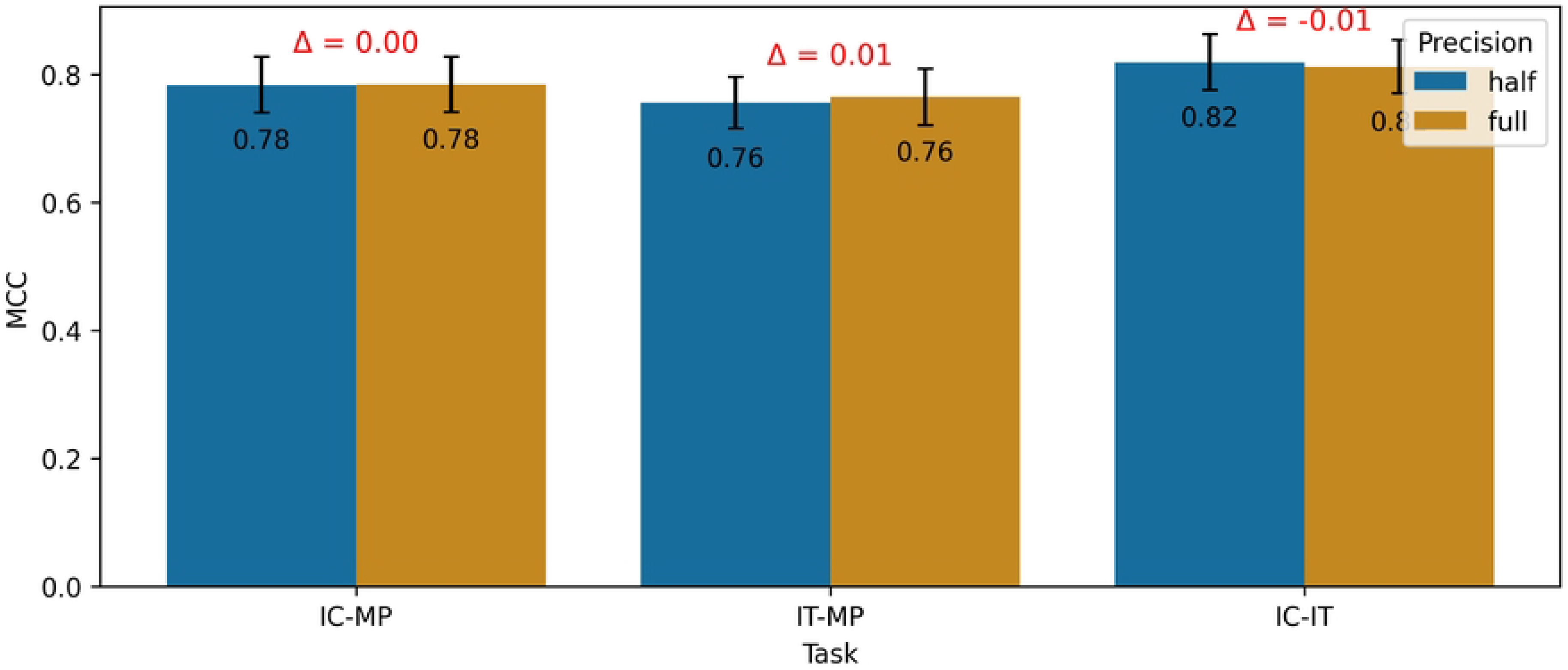

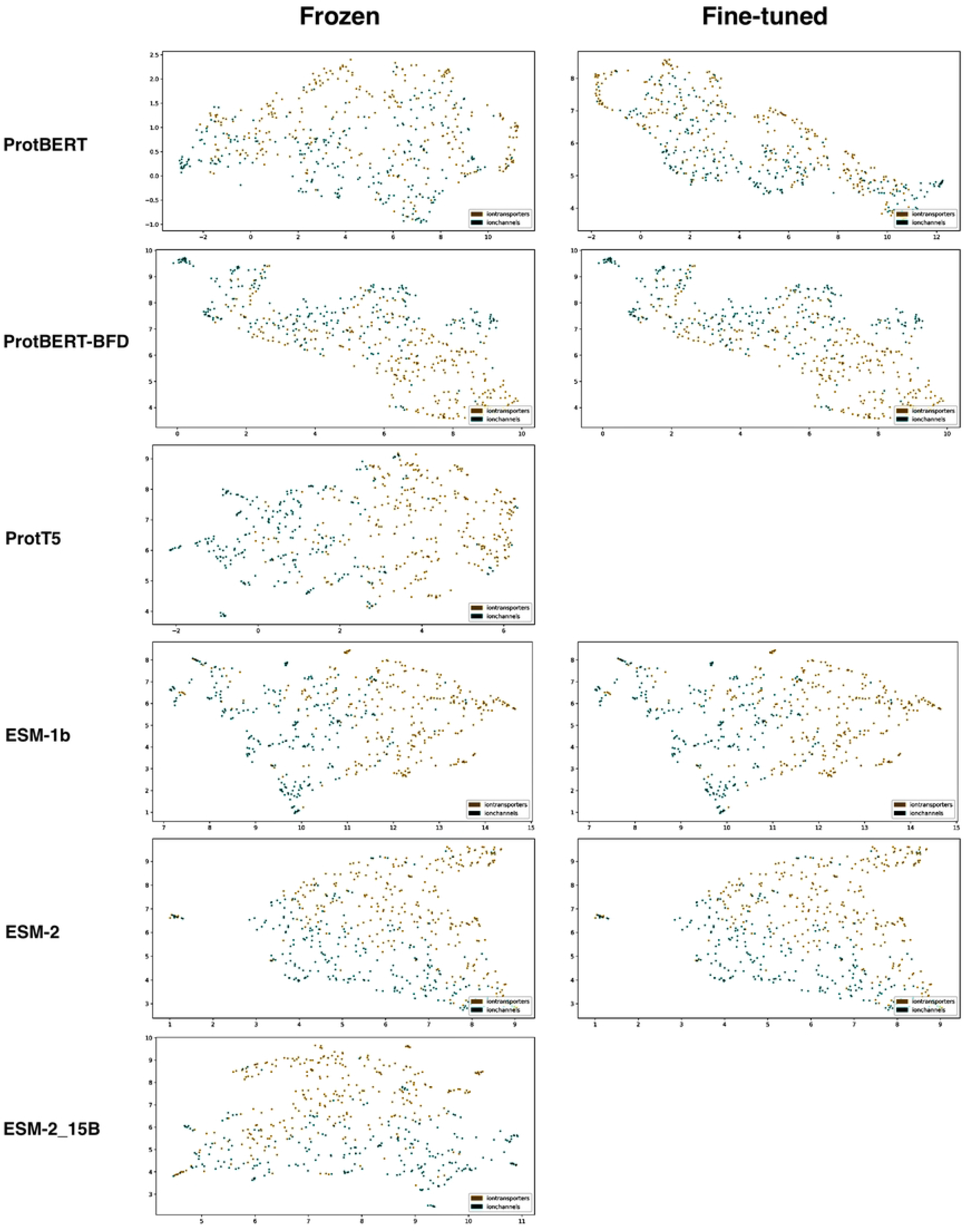

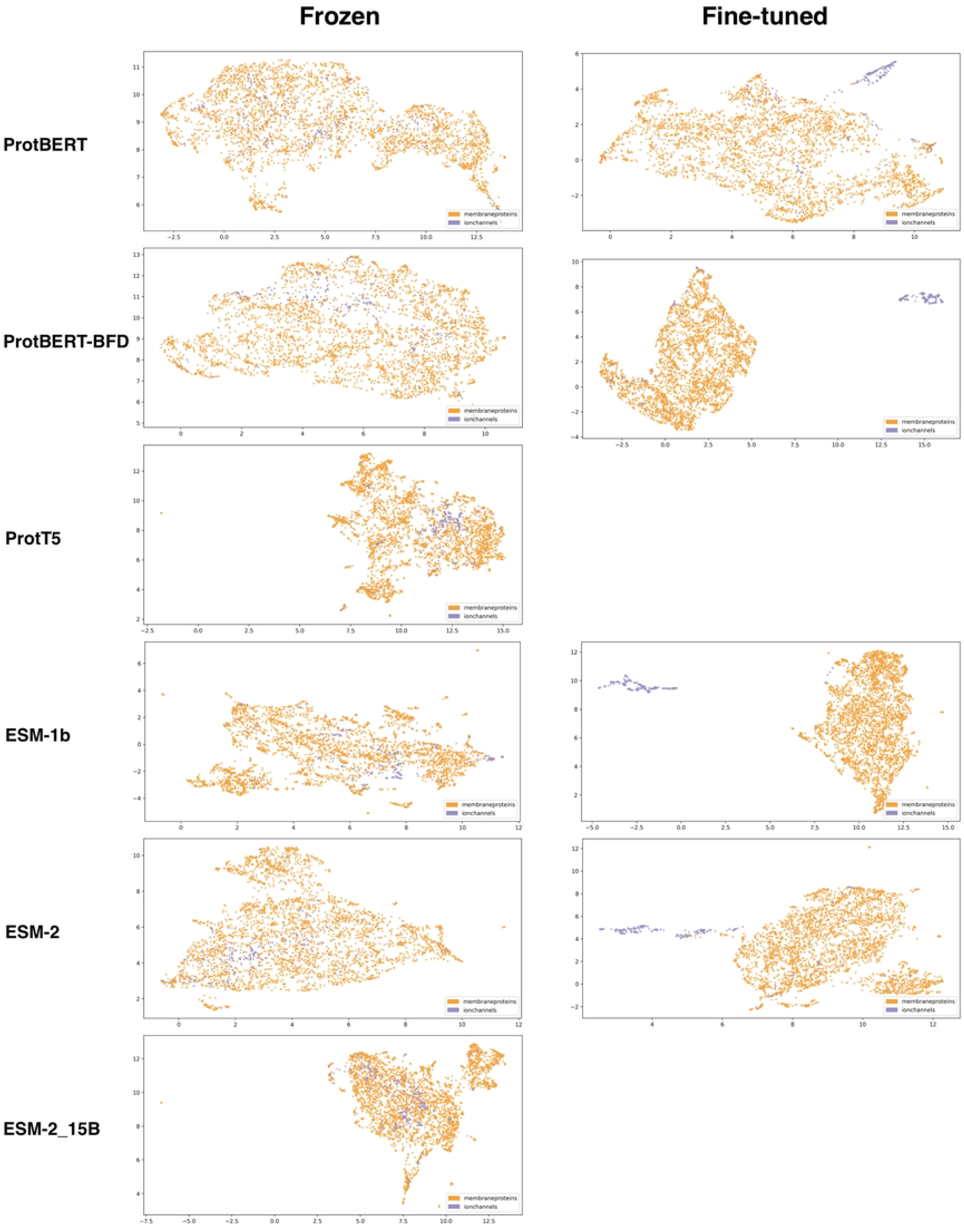

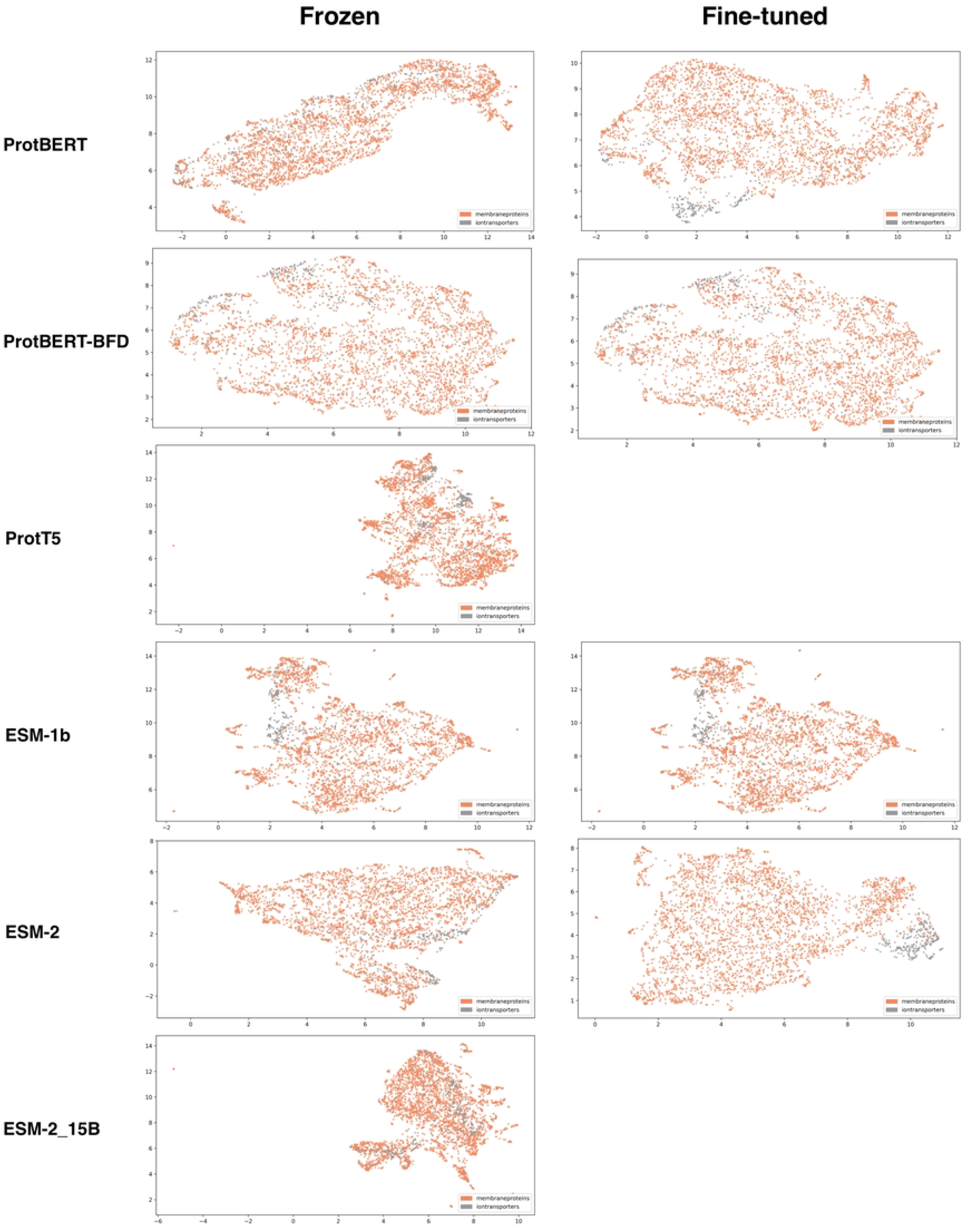

